# Patient iPSC-astrocytes show transcriptional and functional dysregulation in schizophrenia

**DOI:** 10.1101/2020.10.23.350413

**Authors:** Marja Koskuvi, Šárka Lehtonen, Kalevi Trontti, Meike Keuters, Ying Chieh Wu, Hennariikka Koivisto, Anastasia Ludwig, Lidiia Plotnikova, Pekka L. J. Virtanen, Noora Räsänen, Satu Kaipainen, Ida Hyötyläinen, Hiramani Dhungana, Raisa Giniatullina, Ilkka Ojansuu, Olli Vaurio, Tyrone D. Cannon, Jouko Lönnqvist, Sebastian Therman, Jaana Suvisaari, Jaakko Kaprio, Markku Lähteenvuo, Jussi Tohka, Rashid Giniatullin, Claudio Rivera, Iiris Hovatta, Heikki Tanila, Jari Tiihonen, Jari Koistinaho

**Affiliations:** Neuroscience Center, University of Helsinki, Helsinki, Finland; A.I. Virtanen Institute for Molecular Sciences, University of Eastern Finland, Kuopio, Finland; SleepWell Research Program, Faculty of Medicine, University of Helsinki, Helsinki, Finland; Department of Psychology and Logopedics, University of Helsinki, Helsinki, Finland; Department of Forensic Psychiatry, University of Eastern Finland, Niuvanniemi Hospital, Kuopio, Finland; Department of Psychology and Psychiatry, Yale University, New Haven, Connecticut, USA; Mental Health Unit, Department of Public Health Solutions, National Institute for Health and Welfare, Helsinki, Finland; Department of Psychiatry, University of Helsinki, Helsinki, Finland; Institute for Molecular Medicine FIMM, University of Helsinki, Helsinki, Finland; INSERM, Institute of Mediterranean Neurobiology, Aix-Marseille University, Marseille, France; Department of Clinical Neuroscience, Karolinska Institutet, and Center for Psychiatric Research, Stockholm City Council, Stockholm, Sweden

## Abstract

Human astrocytes are multifunctional brain cells and may contribute to the pathophysiology of schizophrenia (SCZ). We differentiated astrocytes from induced pluripotent stem cells of monozygotic twins discordant for SCZ, and found sex-specific gene expression and signaling pathway alterations related particularly to inflammation and synaptic functions. While Ingenuity Pathway Analysis identified SCZ disease and synaptic transmission pathway changes in SCZ astrocytes, the most consistent findings were related to collagen and cell adhesion associated pathways. Neuronal responses to glutamate and GABA differed between astrocytes from control persons, affected twins, and their unaffected co-twins, and were normalized by clozapine treatment. SCZ astrocyte cell transplantation to the mouse forebrain caused gene expression changes in demyelination, synaptic dysfunction and inflammation pathways of mouse brain cells and resulted in behavioral changes in cognitive and olfactory functions. Altogether, our results show that astrocytes contribute to both familial risk and clinical manifestation of SCZ in a sex-specific manner.

## Introduction

Post-mortem studies have revealed reduced brain volume, neuron size, spine density, and abnormal neural distribution in the prefrontal cortex and hippocampus of patients with schizophrenia (SCZ)^1^. Neuropharmacological studies have implicated abnormal dopaminergic, glutamatergic, and GABAergic activity in SCZ^2^, although the molecular mechanisms underlying the disease remain unclear. According to current mainstream theory of the development of the disorder, genetic predisposition to SCZ is pronounced during embryonal development, and environmental effects trigger the symptoms in early adolescence^3^. As post-mortem studies can reveal only findings related to full-blown illness treated with antipsychotics, more information is needed about molecular changes at the early stage of the disease and prior to the psychosis in order to target prevention and early treatment for people in high-risk groups.

Several SCZ-associated genes are involved in the development and physiology of glial cells^4, 5^. Therefore, the importance of astrocytes in neurodevelopmental diseases has gained more interest. Because of their key roles in maintaining CNS homeostasis, synapse formation and the synaptic metabolism of glutamate and monoamines, astrocyte dysfunction may lead to the abnormal neurotransmitter release in SCZ^6^.

Human induced pluripotent stem cell (hiPSC) models have been used to explore neuropathological abnormalities in patients with SCZ, but so far, the main focus has been on neuronal pathophysiology, while the contribution of astrocytes has received much less attention. One study using hiPSC-derived astrocytes obtained from patients with SCZ showed impaired astrocyte maturation related to enhanced BMP signaling pathway in childhood-onset patients with SCZ^7^, and another study found attenuated CCL20 response and T cell recruitment in patients with SCZ after IL-1β exposure^8^. Moreover, transplanted human iPSC-derived glial progenitors obtained from childhood-onset SCZ patients have shown delayed astrocytic differentiation and abnormal astrocyte morphologies as well as behavioral abnormalities in mouse brains^9^.

We previously generated hiPSCs from five monozygotic twin pairs discordant for SCZ and five healthy controls, and showed sex-specific differences in gene expression of hiPSC-derived cortical neurons, especially in pathways associated with N-glycan synthesis, CAMK2G, GABAergic synapse, and purine metabolism in SCZ^10^. In this study, we differentiated hiPSCs from the same twin pairs and healthy controls into astrocytes. Gene expression profiling identified molecules and pathways associated with increased genetic risk for SCZ (between twins and healthy controls) and clinical manifestation of the illness (between affected twins and unaffected twins). We identified sex-specific differences between astrocytes from affected twins, unaffected and healthy controls also previously seen in the patient hiPSC-neuronal dataset, and analyzed in greater depth genes related to synaptic transmission. Furthermore, we demonstrated that astrocytes from SCZ patients dysregulate neuronal calcium responses in astrocyte-neuron co-cultures. We also transplanted our astrocyte progenitors into newborn mouse brains to study behavioral effects and gene expression differences induced by astrocytes from patients with SCZ.

## Results

### Schizophrenia and control hiPSCs differentiate efficiently to astrocytes with similar cell-specific properties

In order to investigate possible astrocyte abnormalities in SCZ^9^, we differentiated our patient hiPSC-lines to astrocytes. The five adult SCZ-discordant monozygotic twin pairs and five healthy unrelated control (Ctrl) hiPSC-lines have been fully characterized before^10^. Three of the pairs were females and two males between the ages of 40 and 69 years (suppl. table 1). Affected female twins (ST) had more severe symptoms than male STs, and they had been treated with clozapine. Males had milder symptoms and used zuclopenthixol, olanzapine, and/or quetiapine. The unaffected twins (HT) had no history of mental health disorders or medication.

We have previously shown that our differentiation protocol generates astrocytes that exhibit stellate-shaped morphology, take up lactate, respond to pro-inflammatory cytokine stimulus, propagate the intercellular calcium waves^11^ and have a gene expression patterns typical of human astrocytes^12^ (Figure 1a). The hiPSC-lines differentiated into astrocytes efficiently regardless of the disease status, and no significant differences were found between the groups or lines in differentiation efficiency based on protein or RNA expression levels of astrocyte specific markers (suppl. figure 1a-b, e-f) and glucose and glutamate uptake (suppl. figure 1c-d).

**Figure 1.**
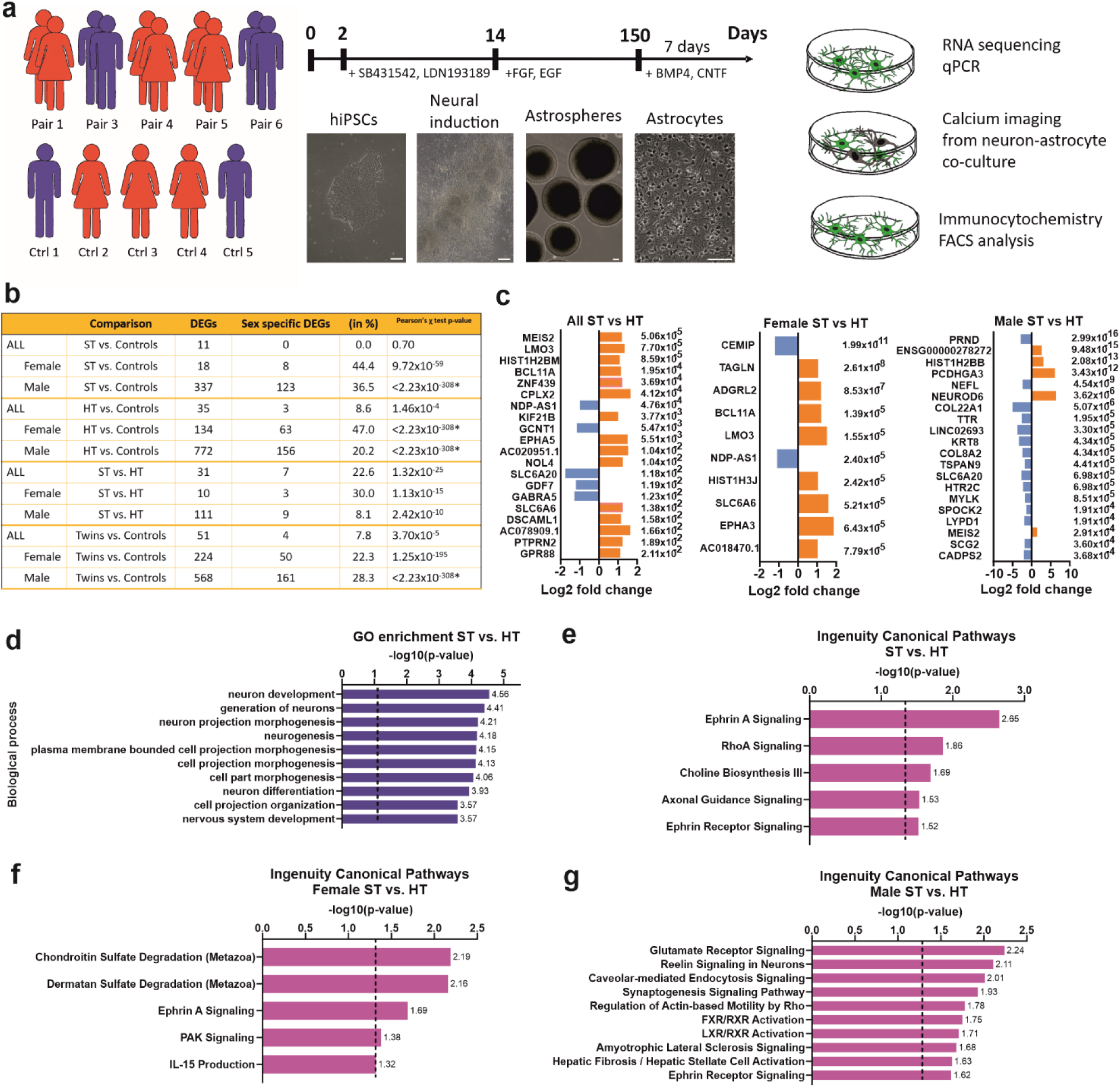
RNA expression analysis of affected (ST) and unaffected (HT) twins using hiPSC-derived astrocytes. **a** The schematic summarizes steps involved in the differentiation of hiPSCs derived from five monozygotic twin pairs discordant to SCZ (red, female; blue, male) and unrelated sex-matched controls towards astrocytes. The protocol mimics embryonal development where astrocyte progenitors arise from neuroprogenitors after being 5-6 months in the culture. Scale bar 50 µm. **b** The comparison table shows the number of differentially expressed genes (DEGs) and proportion of sex-specific DEGs in 12 different comparison sets (DEGs cutoffs: adjusted p-value < 0.05 and log2 change >1.0 in absolute value). Pearson’s Chi-square test p-value indicates the difference in the proportion of sex-specific genes compared with healthy controls having 1.3 % of all genes (461 out of 34,783) differentially expressed between sexes. The p-value for sex-difference was too small to count in the used platform in four comparisons, which reached to 2.23×10-308. **c** The 20 most significant DEGs in ST vs. HT comparison with both sexes, females (containing all 10 DEGs) and males demonstrate the total different gene expression patterns between sexes. **d** The most significant GO enrichment terms in ST vs. HT and ingenuity canonical pathways in ST vs. HT with **e** both sexes, **f** females and **g** males. Dash line is value for p = 0.05. Ctrl = control, ST = affected twin, HT = unaffected co-twin.

### Schizophrenia astrocytes show highly sex-specific gene expression alterations

Sex differences in disease onset, clinical symptoms, and response to drug treatments have been recognized in various psychiatric disorders^13, 14^. We have previously reported sex-specific gene and protein expression differences in hiPSC-derived cortical neurons in patients with SCZ. Our results were also replicable in an independent, larger hiPSC-neuronal cohort^10^. Therefore, we assessed whether sex-specific changes are associated with familial risk or clinical illness of SCZ in hiPSC-derived astrocytes using transcriptomic analysis. Figure 1b summarizes the number of differentially expressed genes (DEGs; Benjamini–Hochberg corrected p-value<0.05 and absolute log2-fold change>1.0) in the comparison between ST and Ctrl (associated with both familial risk and clinical illness), between HT and Ctrl (associated with familial risk without clinical illness), between ST and HT (associated with pure clinical illness) and between all twins and Ctrl (associated with familial risk with or without clinical illness) of all study participants and when separated by sex. While the number of DEGs ranged from 11 between ST and Ctrl to 51 between twins and Ctrl, hundreds of DEGs were revealed in all comparisons when the sexes were analyzed separately. In all comparisons, DEGs were far more numerous in males than females. To find out whether astrocytes contribute to sex differences in the molecular and cellular mechanisms of SCZ, we compared the number of the sex-specific gene expression differences between SCZ astrocytes and Ctrl astrocytes. We first identified sex-specific DEGs among the female Ctrl and male Ctrl astrocytes to compare these genes (1.3%) to the DEGs related to SCZ. Before separating the sexes, the number of sex-specific genes reached up to 22.6 % of the DEGs (ST vs. HT, Pearson’s χ test p =1.32×10^-25^), and after the separation, the proportion was up to 47.0% (Female HT vs. Ctrl, p <2.23×10^-308^). These data highlight sex-specificity of both familial risk and clinical illness-associated changes of gene expression in astrocytes.

The lists of DEGs are found in Supplementary Data 1-12, and the 20 most significant DEGs in the ST vs. HT comparison with combined sexes, females (all DEGs) and males are shown in Figure 1c. Here, the most significant differences were discovered in genes of various functions, such as *BCL11A* (BAF chromatin remodeling complex subunit)^13^, *CPLX2* (Complexin 2)^15^, *SLC6A20* (Solute carrier family 6 member 20)^16^, *GABRA5* (Gamma-Aminobutyric Acid Type A Receptor Subunit Alpha5)^17^, *PTPRN2* (Protein Tyrosine Phosphatase Receptor Type N2)^18^ and *GPR88* (G Protein-Coupled Receptor 88)^19^, all of which have been previously linked to SCZ, and *MEIS2* (Meis Homeobox 2), an autism spectrum disorder-related gene^20^. Notably, in males several collagen genes were differentially expressed. In the ST vs. Ctrl comparison, many of the top genes such as *CNTN1* (Contactin 1), *CDH8* and *CDH13* (Cadherin 8 and 13), *PCDHGB7* (Protocadherin gamma subfamily B, 7), and *CHL1* (cell adhesion molecule L1 like) genes among both males and females were related to cell adhesion. *CHL1* was the number one top gene in the male ST vs. Ctrl comparison (adjusted p =9.6×10^-75^) and, remarkably, also in the male HT vs Ctrl comparison (adjusted p =5.1×10^-93^). We have previously reported *CHL1* to be a top gene in the male ST vs. Ctrl (the second top gene) and the HT vs. Ctrl control (the first top gene) comparisons also in iPSC-derived neurons^10^.

In the HT vs. Ctrl comparison, 5.8 times more DEGs were found in males than in females. The DEGs included downregulation of *SHISA6* (−2.6 log2 fold change, adj. p <0.05), a protein regulating postsynaptic AMPA receptor currents^21^, which was found also in the male comparison (−4.22 log2 fold change, adj. p <0.05). We have previously reported a strong downregulation of this gene in iPSC-derived neurons in the all ST vs. Ctrl and in the male HT vs. Ctrl comparison. In males, also a ribosomal protein gene *RPS4Y1* that has been found to be upregulated in iPSC-derived male neurons in comparisons of the ST vs. Ctrl and the HT vs. Ctrl^10^ was strongly upregulated in the HT vs. Ctrl comparison (13.38 log2 fold change, adj. p < 2.24×10^-17^). In addition, 13 collagen related genes were found in the male comparison. Altogether, these findings indicate that in astrocytes sex is an important determinant of differential gene expression profiles associated with familial risk and clinical illness of SCZ.

### Pathways related to neuronal wiring and inflammation are altered in SCZ astrocytes

We next performed enrichment of DEGs to Gene Ontology (GO) terms and pathways and functional terms of Ingenuity Pathway Analysis (IPA). Total numbers of GO terms (Benjamini-Hochberg, adj. p<0.05) and IPA pathways (−log(p-value)>1.3) in each comparison are listed in supplementary table 2 and the lists of GO terms in Supplementary Data 13-19 and IPA canonical pathways in Supplementary Data 20-28. In general, a greater number of significant GO terms and pathways were detected in males than in females.

In the comparison between ST and HT, 30 GO terms involving neuron development and generation of neurons were detected as the most significant terms (10 most significant GO terms in Figure 1d, all in suppl. data 18). Hypofunction of glutamatergic neurons causing excitation/inhibition imbalance is one of the cellular hypotheses for SCZ^22^. In the male ST vs. HT comparison, Glutamate Receptor Signaling (−log(p)=2.24) was the most significant pathway, and Ephrin receptor pathways known to regulate maturation of synaptic spines and to be linked to SCZ^23^ were altered among both sexes such as Ephrin Receptor Signaling (1.62 in males) and Ephrin A signaling (2.65 combined sexes and 1.69 in females) (Figure 1f-g, suppl. data 27-28).

In addition to Ephrin pathways, neuroinflammation pathways such as IL-15 Production (−log(p)=1.32 in females) and LXR/RXR Activation (1.71 in males) were found in ST vs. HT comparisons. *IL15* (Interleukin-15) is a proinflammatory cytokine, upregulated during neuroinflammation, and it potentially modulates GABA and serotonin transmission causing depression-like behavior, reduced anxiety, and impaired memory^24^. A strongly significant change was in Hepatic Fibrosis Signaling, which was found in the male ST vs. HT comparison, male ST vs. Ctrl and HT vs. Ctrl comparisons, and in the female HT vs. Ctrl comparison. The Hepatic Fibrosis / Hepatic Stellate Cell Activation pathway is the most prominently induced pathway by experimental inflammation in astrocytes^25^ and shares several genes related to extracellular matrix and adhesion with Glycoprotein 6 (GP6) pathway, which was detected in the male ST vs. HT (−log(p)=1.54), ST vs. Ctrl (3.97) and HT vs. Ctrl (6.47) comparisons. Abnormal GP6 and Hepatic Fibrosis / Hepatic Stellate Cell Activation pathways have been previously observed also in iPSC-derived neurons of SCZ in two independent datasets from U.S. and Finland^26^.

In HT vs. Ctrl comparisons, most of the GO terms were related to brain development and synaptic activity. Several GO terms involved MAP kinases in females, and Wnt signaling and collagen related terms in males (suppl. data 16-17). The pathway analysis of the HT vs. Ctrl comparison in females revealed Synaptogenesis Signaling, Ephrin Receptor, and IL-15 Production pathways in addition to Hepatic Fibrosis / Hepatic Stellate Cell Activation pathway (suppl. data 24), whereas in males Wnt, Synaptogenesis, Ephrin, and inflammation related pathways were identified (suppl. data 25).

Together, these results underscore the importance of astrocytes in neuronal synaptic and extracellular processes and inflammation associated with familial risk and clinical illness of SCZ both in males and females.

### The synaptic transmission associated altered gene sets in SCZ astrocytes are distinct in males and females

IPA function and disease pathways in ST vs. HT comparison identified processes related to exocytosis by neurons, motor functions and postsynaptic excitatory processes (Figure 2a-c). The male ST vs. HT comparison showed significant changes in neuritogenesis, brain development and morphology, and schizophrenia disease pathways, whereas in the female ST vs. HT comparison abnormal morphology in brain structures, including development of cerebral cortex and death of its astrocytes, were different. Correspondingly, in the HT vs. Ctrl comparison, pathways of exocytosis/exocytosis by neurons, excitatory postsynaptic processes, and diameter of microvessels were altered, whereas after separation of sexes pathways of (abnormal) morphology and guidance of axons were found both in males and females. Importantly, Schizophrenia disease pathway appeared among the most significant pathways in each male comparison (p=2.14×10^-6^-1.18×10^-7^) (Figure 2d) and was also significant in female twins vs. Ctrl (p=6.93×10^-4^) comparisons. When we looked at the altered genes in each comparison, none of the genes was shared between females and males. However, many of the genes contributing to the schizophrenia disease pathway in astrocytes were related to neurotransmitters and synaptic transmission, and thus also the functional pathway for synaptic transmission appeared significant in separate sex comparisons (Figure 2e).

**Figure 2.**
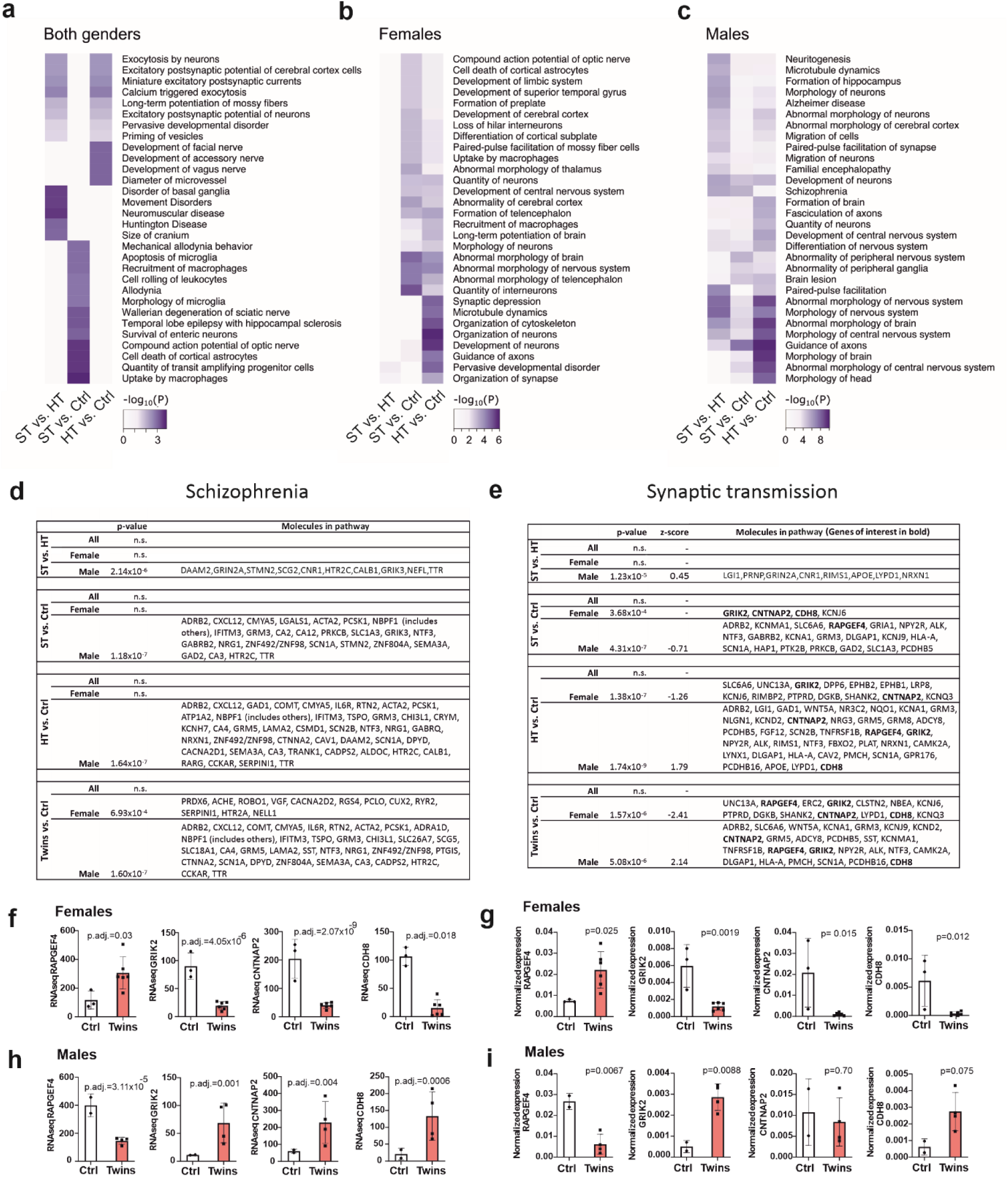
Ingenuity Pathway Analysis of disease and function pathways. The 30 disease and function pathways with the highest p-value in different comparisons among **a** all, **b** female and **c** male groups. The queries are limited to neuronal tissues and CNS cell lines. **d** Schizophrenia disease pathway and related molecules in the pathway in different comparisons. The pathway is categorized as Neurological Disease, Psychological Disorders; schizophrenia; Schizophrenia. **e** Synaptic transmission function pathway and related molecules in the pathway in different comparisons. The pathway is categorized as Cell-To-Cell Signaling and Interaction, Nervous System Development and Function; synaptic transmission; Synaptic transmission. RNA sequence expression of the genes of interest in **f** females and **g** their validation by RT-qPCR and **h** in males and **i** their validation. **f** and **h** have Benjamini–Hochberg corrected p-value, and **g** and **i** have nominal p-value calculated by unpaired t-test. The data is presented as mean ± SD. Ctrl = control, ST = affected twin, HT = unaffected co-twin.

In Synaptic transmission pathway, IPA predicted high decrease for the female Twins vs. Ctrl (z-score= -2.4, p-value= 1.6×10^-6^), but an equally high increase for the male Twins vs. Ctrl (z= 2.1, p= 5.1×10^-6^), respectively. The similar opposite direction in gene expressions between the sexes was also evident between HT and Ctrl comparisons. Among the shared genes with the opposite direction of expression were *RAPGEF4* (Rap Guanine Nucleotide Exchange Factor 4), *GRIK2* (Glutamate Ionotropic Receptor Kainate Type Subunit 2), *CNTNAP2* (Contactin Associated Protein 2), and *CDH8* (Cadherin 8) (Figure 2f-g: females, h-i: males), all of which have been previously associated with SCZ. Also, several voltage-gated potassium channel subunits contributed to an altered synaptic transmission network, but these differed between females (*KCNQ3* and *KCNJ6*) and males (*KCNA1*, *KCNJ9* and *KCND2*). In the ST vs. HT comparison, only males showed significant differences in the synaptic transmission pathway. DEGs included *GRIN2A* (Glutamate Ionotropic Receptor NMDA Type Subunit 2A), *GABRB2* (Gamma-Aminobutyric Acid Type A Receptor Subunit Beta2), *GRM8* (Glutamate Metabotropic Receptor 8), *CNR1* (Cannabinoid Receptor 1), *LYPD1* (LY6/PLAUR Domain Containing 1), and NRXN1 (Neurexin 1). Taking these findings together, our gene expression analysis strengthens the evidence for the involvement of astrocytes in dysregulation of cell adhesion, neuronal wiring, and synaptic functions in a sex-specific manner both in the clinical illness and among those with a familial risk of SCZ.

### Altered pathways in our comparisons were comparable to the published hiPSC-GPC data set with patients with childhood-onset SCZ

For comparison purposes, we performed IPA analysis from DEGs generated in the Windrem et al. data^9^. This previously published RNA sequencing study contained *CD140a^+^* sorted hiPSC-glial progenitor cells (GPCs) derived from 4 patients with childhood-onset SCZ. In total, 116 DEGs between SCZ and control survived the set thresholds in Windrem data set (absolute log2fold change >1.0, FDR 5%, suppl. data 29). The most significant pathways in this data set were Glutamate Receptor Signaling (−log(p)= 4.06), Amyotrophic Lateral Sclerosis Signaling (3.17) and Synaptic Long Term Depression (2.12) (Suppl. figure 2a, suppl. data 30). The first two pathways were shared in our male ST vs. HT and Windrem comparisons together with Synaptogenesis Signaling Pathway (−log(p)= 1.41 in Windrem and 1.93 in male ST vs. HT).

Schizophrenia pathway did not reach significance in this comparison, nor in our pooled sex comparison. Synaptic transmission pathway was significant (p= 1.06×10^-9^) with 15 genes (*FGF12, EPHB1, CSPG5, SHC3, KCNJ9, MPZ, SLC8A3, GRIK4, KCND2, GRID2, FGF14, CHRNA4, GABRA3, SLC6A1*, and *GRIA4*). Four genes (*FGF12, EPHB1, KCNJ9*, and *KCND2*) in four comparisons were shared with the Windrem data set (suppl. table 3). These data indicate that hiPSC-GPCs and hiPSC-astrocytes differentiated in independent studies from childhood SCZ and adulthood SCZ, respectively, share abnormalities in synaptic transmission.

### SCZ astrocytes modulate neurotransmitter responses

As transcriptomics from astrocytes revealed alterations in several pathways related directly or indirectly to synaptic transmission and neurotransmitter functions in SCZ astrocytes, we studied whether ST or HT astrocytes influence neuronal functions. Thus, healthy female hiPSC-cortical neurons were co-cultured with hiPSC-astrocytes from ST, HT, and Ctrl persons. After five weeks in co-culture, astrocytes were positive for astrocytic markers *GFAP*, *AQP4*, and *S100β* and had distinctive stellate morphology (Figure 3a, top panels). Our co-culture neurons were mostly *VGLUT1^+^* glutamatergic neurons and a few were *GABA^+^* GABAergic neurons having embryonal excitatory actions (figure 3a, suppl. figure 3a-b). The responses to glutamate (with co-agonist glycine) and GABA in hiPSC-neurons as calcium influx were recorded (Figure 3b-c).

**Figure 3.**
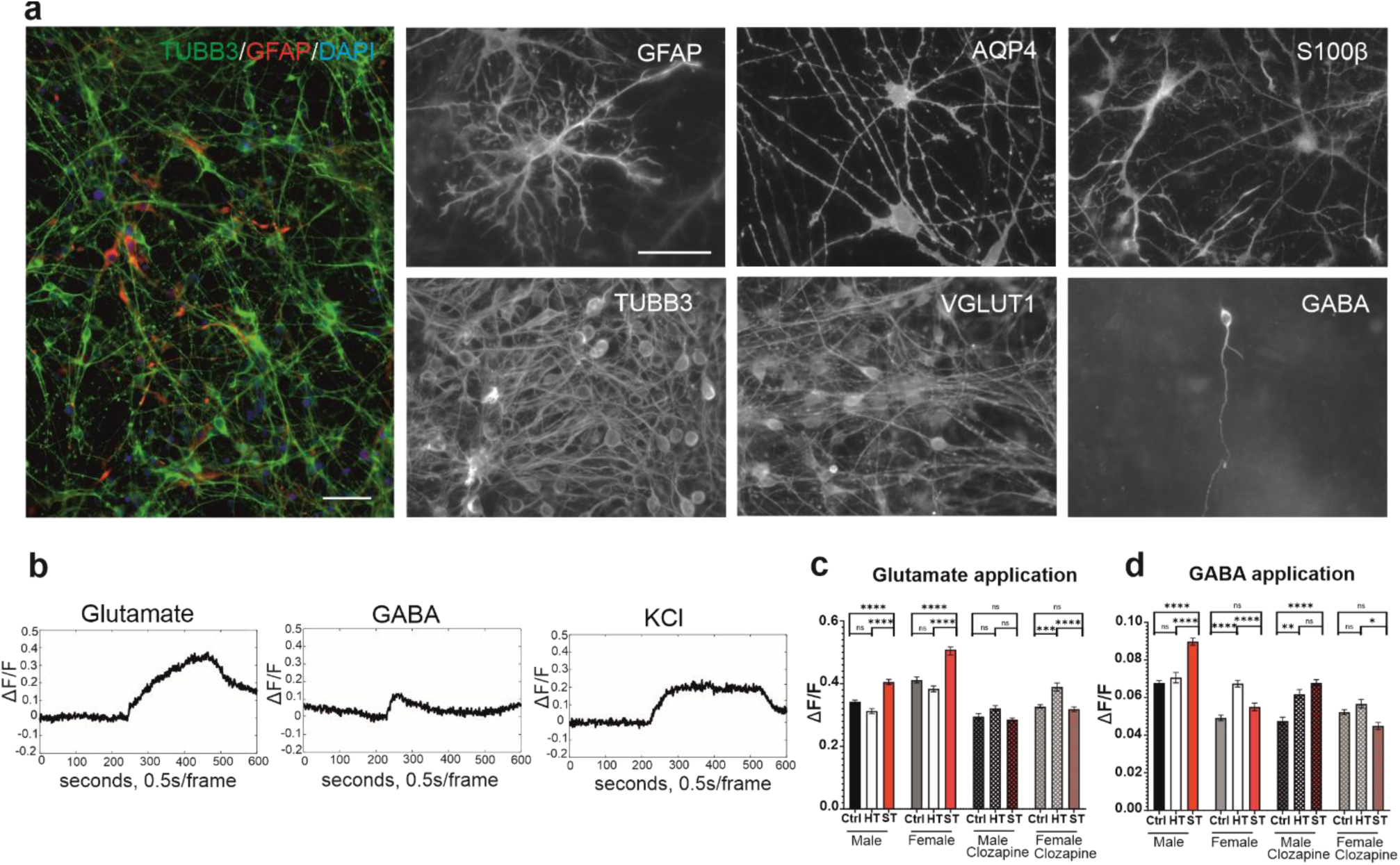
Calcium imaging of control neurons co-cultured with astrocytes. hiPSC-derived astrocytes were plated together with control hiPSC-derived cortical neurons. **a** After 5 weeks in co-culture astrocytes expressed GFAP (red) and neurons TUBB3 (green). Furthermore, astrocyte-specific markers AQP4 and S100β were expressed in these cultures and astrocytes showed a typical stellate morphology. TUBB3+ neuronal cells were positive to VGLUT1 or GABA. The scale bar 50 µm. We used KCl to separate neuronal and astrocytic responses from each other. **b** Examples of calcium traces in neuronal KCl+ cells after application of glutamate with glycine, GABA and KCl. Neuronal responses were presented as ΔF/F values from **d** glutamate and **e** GABA applications. Each group has pooled ΔF/F values from cells from 2-3 subjects, 4-6 recordings, 600-2700 cells per condition. The data is presented as mean ± SD. **** p < 0.0001, ***p < 0.001, ** p < 0.01, 3-way ANOVA and Tukey’s post-hoc test. GFAP = Glial fibrillary acidic protein, TUBB3 = Tubulin Beta 3 Class III, AQP4 = Aquaporin 4, S100β = S100 calcium-binding protein B,VGLUT1 = Vesicular glutamate transporter 1, GABA = Gamma-aminobutyric acid. Ctrl = control, ST = affected twin, HT = unaffected co-twin.

hiPSC-neurons cultured with female Ctrl astrocytes showed a higher glutamate and lower GABA response amplitude (ΔF/F) than those cultured with male Ctrl astrocytes (Tukey post-hoc test, adj. p<0.0001) (Figure 3 d-e, suppl. table 4-5). When the effect of astrocytes derived from twins were compared to astrocytes derived from same-sex Ctrl, no significant differences were detected in glutamate response between HT and Ctrl, whereas neurons co-cultured with ST astrocytes of either sex showed significantly increased response to glutamate (adj. p<0.0001) when compared to neurons co-cultured with HT or Ctrl astrocytes. Neurons with male ST astrocytes showed a significant increase in calcium response after GABA treatment (adj. p<0.0001) compared to neurons with Ctrl and HT astrocytes, whereas in females neurons co-cultured with female HT astrocytes showed an increased response (adj.p<0.0001) compared to control and ST astrocytes. The proportions of responding neurons were similar between the astrocyte groups (suppl. figure 3c-d). The data indicate that regulation of neuronal responses to neurotransmitters is differentially altered in male and female astrocytes derived from individuals with clinical illness and familial risk of SCZ.

We have previously shown that clozapine treatment normalizes the calcium responses in ST hiPSC-neurons^10^. After clozapine application to the co-cultures, the increased calcium response to glutamate observed in neurons co-cultured with ST astrocytes decreased (adj. p<0.0001) to Ctrl level in both the males and females compared, but the treatment was inefficient in female HT astrocyte co-cultured neurons, leaving their calcium responses higher than in female ST and Ctrl astrocyte co-cultured neurons. When GABA was applied to clozapine-treated cells, the calcium response decreased differentially in each group, leaving it significantly increased in males both in ST (adj. p<0.0001) and HT (adj. p<0.01) astrocyte co-cultured neurons, whereas in females, the responses in HT astrocyte co-cultured neurons was higher (adj. p<0.05) than in ST astrocyte co-cultured neurons after GABA application. These results link abnormal expression of synaptic transmission genes in astrocytes with altered regulation of neuronal transmitter responses by ST and HT astrocytes to different effects of antipsychotic treatment in a sex-specific manner.

### Transplanted hiPSC-astrocyte progenitors mature into astrocytes and induce subtle behavioral changes

We next asked whether human SCZ astrocytes affect neuronal gene expression or brain functions in *in vivo*. For this purpose, we humanized mice by transplanting female ST, HT, and Ctrl hiPSC-astrocyte progenitors into neonatal mouse forebrains and analyzed mouse behavior at five and 10 months followed by gene expression profiling of the frontal cortices seven days later (Figure 4a, for characterization, see Methods, figure 4b-d, suppl. figure 4). A behavioral test battery covering motor, sensory, emotional, social and cognitive functions relevant for SCZ was used (for more details, see Methods). In a Novel Object Recognition (NOR) test experimental objects caused high stress so that 38.1% of ST mice compared to 12.5% and 22.2% of Ctrl mice and HT mice, respectively, were disqualified from the test at five months (suppl. table 6). However, the NOR test of the remaining mice showed a significant age x transplantation interaction (p = 0.02, General linear model, repeated measures), such that ST mice had preference for novelty as they aged (Figure 4e). Also, ST mice were more interested in the cardamom odor compared to the other groups (preference time p=0.007), which might indicate increased sensitivity to odors. Of note, patients with SCZ have been reported to have olfactory dysfunctions^27^. The other behavior tests did not identify any significant differences between the groups (Suppl. figure 5). Thus, neonatal transplantation of SCZ astrocyte progenitors resulted in mild behavioral changes associated with cognitive and olfactory functions by the age of 10 months.

**Figure 4.**
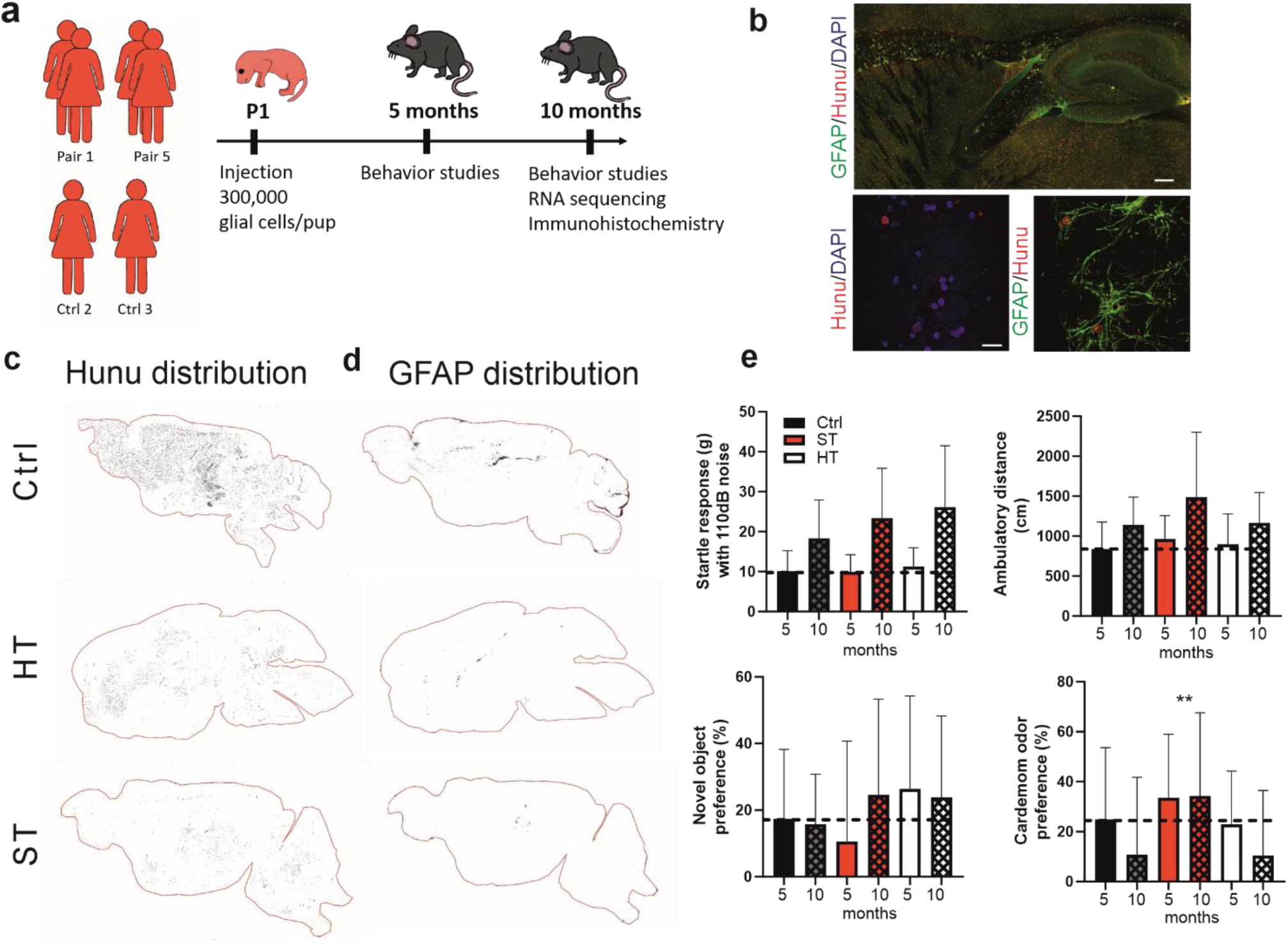
Transplantation of human astrocytes into neonatal mouse brain. **a** Schematic illustration of hiPSC-glial progenitor chimeras that were established by neonatal injections into Rag 1 KO hosts and sacrificied after 10 months. **b** Representative pictures from Hunu^+^ cells (human nuclei) and human GFAP^+^ cells. Sagittal sections demonstrating the distribution of **c** Hunu and **d** GFAP. A portion of the transplanted cells expressed human-specific GFAP and the highest density of the GFAP^+^ human astrocytes were found in the corpus callosum. **e** Results from behavioral studies: Novel object preference, ambulatory distance, Pre-pulse inhibition with a 50 dB tone and cardamom odor preference of the transplanted mice. General linear model, repeated measures and Tukey’s post-hoc test. ** p < 0.01. Ctrl, n=19, ST n=24, HT n=27. The results are presented as mean ±SD. Ctrl = control, ST = affected twin, HT = unaffected co-twin.

### Transplanted SCZ astrocytes induce demyelination, synaptic dysfunction and inflammation pathways in mouse brain cells

We chose to use frontal cortical tissues of the humanized mice for transcriptomic analysis, as this brain area shows early involvement in SCZ pathophysiology^28^. The analysis was done with mouse cell-derived reads (for more details, see Methods, suppl. figure 6a-b, and suppl. data 31). In this transcriptomic analysis the threshold of 0.05 for nominal p-value was used as DEG criteria as only few of the mouse DEGs survived Benjamini-Hochberg adjustment (suppl. table 7). In ST mice vs. HT mice or Ctrl mice, GO terms identified significant biological processes related to nervous system development and myelination (Figure 5a-b), and included many of the most significant downregulated myelination-related DEGs such as *Cldn11* (Claudin 11), *Mobp* (Myelin-associated oligodendrocyte basic protein), *Mbp* (Myelin basic protein), *Plp1* (Proteolipid protein 1) and *Tspan2* (Tetraspanin 2) (figure 5d, suppl.figure 6c-d, suppl.data 32-34).

**Figure 5.**
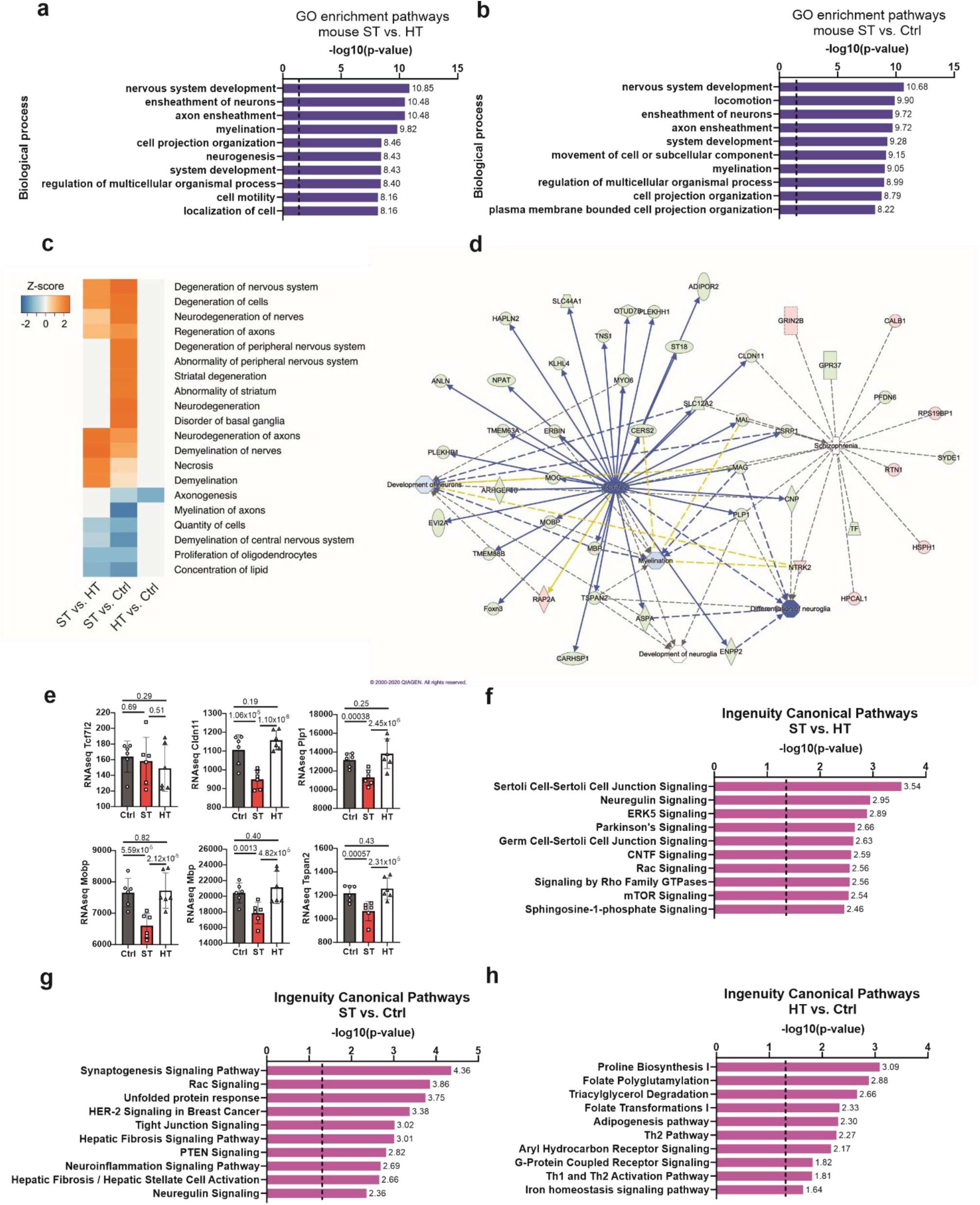
RNA sequencing analysis from mouse brain after transplantation. The first 10 GO enrichment pathways between **a** ST and HT mice and **b** ST and Ctrl mice. **c** The 20 disease and function pathways with the highest z-scores in different comparisons. **d** Functional network maps for TCF7L2 regulated genes and their association to myelination, development of neurons, differentiation of neuroglia and schizophrenia. Red indicates up-regulated genes and green down-regulated genes in color intensity-dependent manner. **e** RNAseq expression of mouse Tcf7l2 and the regulated genes regarding myelination. All data are presented as mean ± SD. The first 10 canonical pathways between **f** ST and HT, **g** ST and Ctrl and **h** HT and Ctrl. n = 6 for ST, HT and Ctrl mice, dash line is value for p = 0.05. Ctrl = control, ST = affected twin, HT = unaffected co-twin.

The full lists of GO terms are in Supplementary Data 35-37 and IPA canonical pathways in Supplementary Data 38-40. When top z-scores in disease and function pathways were combined in different comparisons (Figure 5c), pathways of demyelination, a previously implicated hallmark of SCZ, and neurodegeneration were identified in ST mice vs. HT or Ctrl mice. Based on human specific immunostainings (figure 4 b, suppl. figure 4) and RNA expression profile of human-derived reads (suppl. figure 6b), majority of the transplanted glial cells maturated to astrocytes (for more details, see Methods). As the analyzed sequence reads are of mouse origin and astrocytes are non-myelinating cells, this could point to astrocyte-regulated oligodendrocyte dysfunction in ST and even in HT mice. Also proliferation of oligodendrocytes –pathway was downregulated in ST mice vs. HT mice and in ST mice vs. Ctrl mice. In comparison of ST mice vs. HT or Ctrl mice, we identified several canonical pathways related to synaptic functions that have been previously associated with SCZ, such as Neuregulin (−log10(p)=2.36) and Rac signaling (3.86) in ST mice vs. Ctrl mice, and Neuregulin (−log10(p)=2.95), ERK-5 (2.89), Rac Signaling (2.56) and Signaling by Rho family GTPases (2.56) in ST vs. HT mice (Figure 5f-h). In addition, synaptic pathways signaling was strongly affected in the ST vs. Ctrl comparison. Importantly, in ST vs. Ctrl mice we detected major pathways related to inflammation, such as Hepatic Fibrosis Signaling (−log10(p)=3.01), Hepatic Fibrosis / Hepatic Stellate Cell Activation (2.66) and Neuroinflammation Signaling (2.69), and in ST mice vs. HT mice Neuregulin Signaling and Hepatic Fibrosis Signaling Pathways (−log10(p)=1.67), and in HT mice vs. Ctrl mice Neuroinflammation Signaling Pathway (−log10(p)=1.59).

Finally, the causal pathways comparing ST vs. HT identified *SH3TC2*, *TARDBP*, and *TCF7L2* as regulators of astrocytosis, demyelination of nerves, and neurodegeneration of axons (consistency score = +7.54). Here, transcription factor 7-like 2 (*TCF7L2*) was the main master regulator of hypomyelination and neurodegeneration detected in the ST vs. HT comparison (p-value of overlap = 1.35×10^-35^, activation= -7.14) or ST vs. Ctrl comparison (p-value of overlap= 2.22×10^-17^, activation= -5.43) (suppl. data 41-43). Inhibited Tcf7l2 mediated gene regulation connects many of the significant DEGs and pathways under one regulator (figure 5d-e). Altogether, these results indicate that human ST astrocytes alter mouse brain gene expression to promote demyelination, neuroinflammation and altered synaptic function, all features of SCZ neuropathology.

## Discussion

The importance of astrocytes in neurodevelopment and disease has been increasingly appreciated because of their key roles in maintaining central nervous system homeostasis and their close interactions with neurons, other glial cells (microglia and oligodendrocytes), and blood vessels in the brain, but their role in neurodevelopmental disorders is not well-known.

Previous hiPSC-astrocyte studies have addressed the defects in SCZ related to differentiation, maturation, or inflammatory responses. The cultured hiPSC-astrocytes from monozygotic twin pairs discordant for SCZ replicated some of the previous findings in SCZ-derived hiPSC-astrocytes. As in Akkouh, et al.^8^, our hiPSC-astrocytes derived from patients with SCZ had no significant differences in their differentiation efficacy, basic astrocytic marker expression, or glutamate uptake. Based on the transcriptomic data, cultured astrocytes of affected twins showed altered pathways related to inflammation, while SCZ-derived hiPSC-astrocytes in the study by Akkouh et al. responded differentially to IL1β-induced inflammation. Liu, et al.^7^ reported that astrocyte differentiation and maturation are impaired in glial progenitor cells derived from childhood-onset SCZ. Similar to our study, they found a number of potassium-related genes to be differentially expressed in SCZ glial cells, and further detected reduced uptake of potassium in these cells. These previous studies involved patients from both sexes, but did not address sex differences in SCZ.

Astrocytes are in close contact with neuronal synaptic clefts for receiving synaptic information from released neurotransmitters and their quantities^29^. Our data showed aberrant expression of glutamatergic and GABAergic receptor genes in SCZ astrocytes. Especially *GRIK2* was sex-specifically expressed in astrocytes of affected and unaffected twins, being significantly upregulated in males, but downregulated in females when compared to same-sex healthy controls. Similar sex-specific disruptions have been observed in the molecules regulating glutamate receptors in the post-mortem dorsolateral prefrontal cortex of patients with major depressive disorder^30^. Furthermore, the glutamate receptor signaling appeared in the male but not female affected vs. unaffected twin comparison. These data suggest that the excitatory/inhibitory imbalance in SCZ may be due to the primary dysfunction of the glutamatergic or the GABAergic signaling pathway and that the previously hypothesized NMDA receptor hypofunction in glutamatergic neurons as a major defect in SCZ^31^ may actually be a secondary change.

Astrocytic contribution to excitatory/inhibitory imbalance is likely related to reduced gliotransmitter release and/or insufficient clearance of neurotransmitters^32^. Clozapine activates astroglial release of L-glutamate and D-serine^33^, and reduces glutamate uptake dose-dependently in rodent cell cultures^34^. We saw an increase in the neuronal responses to glutamate and GABA application as elevated internal calcium signal in healthy neurons co-cultured with affected twin astrocytes. Clozapine was able to reduce the responses in affected twin astrocyte co-culture almost to the same level as in healthy control co-cultures. Our previous calcium influx study on patient hiPSC-neuronal cultures without astrocytes indicated an increased response to glutamate treatment in female affected twins and a decreased response to GABA in male affected twin compared to unaffected twin neurons, while clozapine normalized the differences to a non-significant level^10^. Even though the analysis protocols differ, we saw more disease-related differences in affected twin astrocyte co-cultured neurons than previously in affected twin neurons.

Among the extracellular matrix (ECM) proteins, cadherin involvement is evident in major psychiatric disorders^35^, and aberrant expression of collagens in depressive and aggressive male mice indicates ECM remodeling to occur in different psychopathological states^36^. Our identification of a large number of adhesion and collagen genes that are differentially expressed in astrocytes between affected and unaffected twins, and between unaffected and healthy individuals, indicates involvement of ECM in SCZ. Notably, we demonstrated strongly altered astrocyte expression of neural cell adhesion molecule L1-like protein (*CHL1*) between affected and unaffected males and females, and even between unaffected and healthy males. *CHL1* regulates neuronal survival and growth, and functions together with Semaphorin 3B to induce dendritic spine pruning in developing pyramidal neurons^37^. *CHL1* has been linked to SCZ in several studies and its expression is regulated by clozapine^38^. It was also one of the most altered genes by expression in the male affected twins vs. healthy controls and unaffected twins and healthy controls comparisons in iPSC-derived neurons^10^. Abnormalities in collagen genes are associated with SCZ^25^ and in animal models loss of certain collagen molecules results in decreased inhibitory synapse number, reduction in perineuronal nets, and SCZ-related behaviours^39, 40^. A large body of literature shows abnormalities in perineuronal nets and ECM in SCZ^41, 42^, and implicates ECM in regulation of cell migration and neurite outgrowth^43^, and perineuronal nets in protection of parvalbumin-expressing GABA interneurons^42^. Taken together, we identified links between altered gene expression in inflammation, ECM, and synaptic pathways in astrocytes of individuals with clinical illness and familial risk, which are associated with altered regulation of neurotransmitter responses in neurons.

To our knowledge, only one hiPSC-glial transplantation study including patients with SCZ has been published^9^. Transplanted hiPSC-glial progenitors from a patient with childhood-onset SCZ had aberrant migration, reduced myelination and abnormal astrocytic morphologies in the corpus callosum in shiverer mice. Behavioral deficits in various tests were found. As the symptoms of childhood-onset SCZ appear more than a decade earlier and are far more severe than in SCZ, we may not have seen clear symptoms in our behavioral tests because the mice were still in a prodromal stage or simply due to the less severe phenotype of SCZ compared to childhood-onset SCZ or different differentiation methods of the cell lines.

Patient studies have implied impaired myelination, accelerated brain aging, neurodegeneration, upregulated brain immune responses, and neuroinflammation in SCZ subjects^44, 45^. *TCF7L2*, which was the main regulator of hypomyelination and neurodegeneration in mice transplanted with astrocytes derived from SCZ patients, encodes a member of the LEF1/TCF family of transcription factors that cooperates with β-catenin in the canonical Wnt signalling pathway, the key pathway controlling both oligodendrocyte development and myelination processes^46^. *TCF7L2* polymorphisms have been associated with bipolar disorder, SCZ, and autism. Moreover, mice with modulated *TCF7L2* expression (either null or overexpressed) have shown differences in fear learning, but not on the PPI test^47^. Our IPA results also demonstrated that lenalidomide and lithium could increase the expression of *TCF7L2*-related pathways. Lenalidomide is used to treat multiple myeloma and myelodysplastic syndromes, and it has been tested to improve autism symptoms and could potentially reduce cytokine inflammation^48^. Lithium is a mood stabilizer, commonly used to treat bipolar disorder and major depression. Due to its neuroprotective properties in cells, astrocytes are potential direct targets for lithium treatment^49^.

In conclusion, our results showed that signaling pathways related to inflammation, synaptic functions, and especially, collagen and glycoprotein 6 pathways contributing to ECM are crucial in the etiology of SCZ. Abnormalities in ECM have been among the strongest findings in iPSC-derived neurons in 3 other datasets^26, 50^, which highlights its importance in the pathophysiological process. Finally, our results emphasize the importance of analyzing gene expression results separately for males and females. The reproducible results suggest the intriguing possibility that SCZ may be primarily a disease of ECM of the brain: abnormal and insufficient cell adhesion and ECM may result into abnormal neuron guidance and synaptogenesis, leading to imbalance in GABA/glutamate balance, and, finally, to clinical symptoms.

## Methods

### Patient iPSC-lines

The Ethics Committee of the Helsinki University Hospital District has approved this project (statement no. 262/EO/06). Five monozygotic twin pairs discordant for SZ and five age- and sex-matched healthy volunteers were included. Patient fibroblasts were obtained from skin biopsies and reprogrammed to hiPSCs with CytoTune-iPS 2.0 Sendai Reprogramming Kit (ThermoFisher Scientific). The patients’ descriptions and full characterization of patients and control iPSC lines are published in Tiihonen et al. 2019^10^.

### Astrocyte differentiation

The hiPSC-derived astrocytes were differentiated based on the previously published protocol^51^ with slight modifications. The hiPSCs were grown on Matrigel-coated (BD) dishes in E8 medium (Gibgo). The medium was changed every other day, and hiPSC-colonies were enzymatically passaged by 0.5 mM EDTA weekly (Gibgo). hiPS-cells were differentiated to neuroepithelial cells by dual SMAD inhibition: using SB431542 10 µM (Sigma) and LDN-193189 200 nM (Miltenyi Biotec) for 12 days as previously described in Oksanen et al. 2017^11^. The cells were detached from Matrigel-coated plates to ultralow attachment dishes in astrocyte sphere medium (ASM) containing DMEM/F12 medium supplemented with 5 % N2 supplement, 2 mM Glutamax, 50 IU/ml penicillin, and 50 μg/ml streptomycin (all from Gibgo) and 5000 KY/ml Heparin (LEO), 25 ng/ml bFGF and 25 ng/ml EGF (both R&D Systems). Half of the medium was renewed every 2-3 days, and spheres were manually cut once a week. At the age of 5 months, astrocyte progenitors were dissociated with Accutase (Stemcell technologies), plated as 100k/well on Matrigel-coated 24-well plates in ASM medium and maturated with 10 ng/ml CNTF and 10 ng/ml BMP4 for one week.

### Neuron-astrocyte co-culture

We used the same protocol described in Oksanen et al. 2017^11^. One control hiPSC-neuron line (control 3) was co-cultured with hiPSC-derived astrocytes obtained from all SCZ pairs. The neuro- and astro-spheres were dissociated to single cells with Accutase and mixed at a ratio of 1:0.75, hiPSC-derived neurons and hiPSC-derived astrocytes, respectively. The diluted Matrigel at 1:6 was used for embedding. Totally around 50,000 cells were plated as thin-layer culture onto poly-L-ornithine coated circular cover glasses (9 mm of diameter, Thermo Fisher Scientific). Co-cultures were cultured 5 weeks in neurosphere medium (NSM): 1:1 DMEM/F12 and Neurobasal supplemented with 5% N2 supplement, 2 mM Glutamax and 50 IU/ml penicillin, and 50 μg/ml streptomycin (all from Gibco).

### Glucose uptake

hiPSC-derived astrocytes were plated on a 12-well plate at 300k/well for one week in maturation medium. The cells were incubated in the presence of 120 µM 2-NBDG (2-Deoxy-2-[(7-nitro-2,1,3-benzoxadiazol-4-yl)amino]-D-glucose, Thermo Fisher Scientific) for 30 minutes in the glucose-free medium. The cells were collected and resuspended in PBS. Flow cytometry (FACS Calibur) counted the number of cells that had taken-up glucose.

### Glutamate uptake

hiPSC-derived astrocytes (n=2 per group) were plated on a 12-well plate at 300k/well for one week in maturation medium. Cells were washed with BSS solution (3.1 mM KCl, 1.2 mM CaCl_2_, 1.2 mM MgSO_4_, 0.5 mM KH_2_PO_4_, 5 mM PIPES, 2 mM glucose, pH 7.2) containing either 135 mM NaCl or 135 mM C5H14ClNO (Choline chloride), and incubated 10 minutes at 37 °C by addition of 1 μCi/ml of L-[3,4-3H]-Glutamic acid (Perkin Elmer, NET490) and 10µM L-glutamate (Sigma). Cells were washed and lysed. The lysate was mixed with Ultima Gold (Perkin Elmer) and the relative expression of Na^+^-dependent uptake of radiolabeled glutamate was calculated based on the measurement with a microplate scintillation counter (1450 MicroBeta Plus, Wallac).

### Immunocytochemistry

Plated hiPSC-derived astrocytes were fixed in 4% PFA for 20 minutes. The primary antibodies for monolayer astrocyte cultures mouse anti-GFAP 1:800 (Chemicon MAB360) and rabbit anti-S100β, 1:500 (Swant 37a), and for co-cultures mouse anti-TUBB3 1:1000 (Biolegend 801201), rabbit anti-VGLUT1 1:300 (Sigma, V0389-200), rabbit anti-GABA 1:600 (Sigma A2052), rabbit anti-AQP4 1:500 (Sigma-Aldrich AB3594), rabbit anti-GFAP, 1:500 (Dako Z0334) and rabbit anti-S100β 1:100 (Abcam ab52642) were incubated overnight in +4 °C. The secondary antibodies Alexa Fluor goat anti-rabbit 568 (Invitrogen A11008) and Alexa Fluor goat anti-mouse 488 (Invitrogen A11001) (1:300 dilution) were added for 1 hour at room temperature. Nuclei were visualized by DAPI staining (Sigma) for 5 minutes at room temperature. Images were taken using Zeiss AXIO Observer microscope.

For flow cytometric quantification, plated hiPSC-astrocytes were detached and samples for intracellular stainings were fixed in eBioscience Foxp3 fixation solution (Thermo Fisher Scientific) overnight and primary antibodies rabbit anti-GFAP 1:75 (Dako Z0334) and mouse anti-VIM 1:25 (Sigma V2258) were incubated for 30 minutes in +4 °C. The secondary antibodies R-phycoerythrin goat anti-rabbit IgG 1:500 (Invitrogen A10542) and R-phycoerythrin goat anti-mouse IgM 1:400 (Invitrogen, A10689) were added for 30 min at room temperature. Samples for surface staining were incubated with human FcR Blocking Reagent (Miltenyi Biotec) together with PE-conjugated mouse anti-GLAST 1:15 (Miltenyi Biotec 130-095-821) for 15 minutes. At least 15k cells/sample were collected with FACS Calibur.

### Calcium imaging and analysis

hiPSC-derived neuron and astrocyte co-cultures were loaded with Fluo 4 direct (calcium assay kit by Thermo Fisher Scientific) for 30 minutes in 37 °C. During imaging, the cells were continuously perfused with basic solution (152 mM NaCl, 10 mM glucose, 10mM HEPES (pH 7.4), 2.5 mM KCl, 2 mM CaCl_2_ and 1 mM MgCl_2_). The flow was controlled by a peristaltic perfusion system PPS2 (Multichannel systems) to a 3 ml/min flow rate in total volume of 2 ml. The hiPSC-derived neurons were stimulated sequentially for 2 minutes with glutamate (100 µM) together with NMDA receptor co-agonist glycine (10 µM), and then with GABA (100 µM). After that, we used KCl (30 mM) for depolarizing stimulation to separate neuronal and astrocytic responses from each other^52–54^. Ionomycin (10 µM, Cayman) was added at the end of the imaging session to test vitality of cells. The recordings were performed with an inverted Zeiss AXIO Observer microscope equipped with Prime BSI sCMOS camera (Photometrics).

The results were analyzed using a custom script in MatLab software (MATLAB R2019b). Analysis was blind to experimental conditions and sample identity. Regions of interests, corresponding to individual cells, were masked and the mean pixel intensity was measured at each time point. Traces were detrended and intensity of the treatment period was normalized to the baseline intensity to produce ΔF/F.

### RT-qPCR

RNA was extracted from the monolayer culture of hiPSC-derived astrocytes using the RNeasy Mini kit (Qiagen) according to the manufacturer’s protocol. The extracted RNA was DNAse-treated and eluted in nuclease-free water. RNA from transplanted mice was extracted using Trizol (Sigma) phase separation and purified with the RNeasy Mini kit. Synthesis of cDNA was performed using Maxima reverse transcriptase enzyme approach (Thermo Fisher Scientific). Maxima Probe qPCR Master Mix (Thermo Fisher Scientific) and Taqman assay from Thermo Fisher Scientific were used. Human primers used were: CDH8 (Hs00242416_m1), CNTNAP2 (Hs01034296_m1), RAPGEF4 (Hs00199754_m1), GRIK2 (Hs00222637_m1), and mouse primers: Cldn11 (Mm00500915_m1), Plp1 (Mm01297210_m1), Mobp (Mm02745649_m1), Mbp (Mm01266402_m1), and Tspan2 (Mm00510514_m1). The results were normalized to ACTB (Applied Biosystems) in human gene expression and to B2m (Mm00437762_m1) in mouse using Q-gene program (Equation 2)^55^.

### RNA sequencing

RNA-sequencing libraries were produced with NEBNext Ultra II Directional RNA Library Prep Kit (New England Biolabs) from hiPSC-derived astrocytes of ST, HT and Ctrl (all groups n=5; 2 males and 3 females), and from cortices of 10-month old mice that received transplanted hiPSC-derived astrocytes from ST, HT or Ctrl subjects (altogether 18 mice cortices and 9 hiPCS-derived astrocytes; n= 6 mice/group, n= 3 hiPSC-derived astrocytes/group), as well as three cortices from vehicle controls with no transplanted cells. Libraries were sequenced with NextSeq500 (Illumina, hiPSCs average 41.0 and transplanted cortexes average 39.2 million reads per sample). The sequencing was provided by the Biomedicum Functional Genomics Unit at the Helsinki Institute of Life Science and Biocenter Finland at the University of Helsinki. For hiPCS-derived astrocytes, sequence reads were aligned to the human genome GRCh.38 using STAR aligner v2.7.2^56^ and annotated to gene exons with HTSeq v0.11.2^57^. For transplanted samples, the sequence reads were aligned simultaneously to the human and mouse (GRCm.38) genomes with STAR, allowing identification and separation of sequences that originated from mouse or human cells, based on nucleotide variation between the species, and were annotated to gene exons as above. On average, 90.3% of total reads had a unique best alignment on either genome and were usable for HTSeq annotation. Alignments of the non-transplanted controls showed that, on average, less than 0.03% of sequences (783 protein coding genes) misaligned between species (Supp. Data 31). The number of reads aligned with human genome were significantly less in ST and HT mice compared to Ctrl mice (suppl. figure 5a). To validate that the human-aligned sequence reads match with astrocytes, we did a deconvolution analysis using MuSiC^58^ with a brain derived single-cell sequencing data^59^ as cell type reference. The transplanted cells clustered 93 % to astrocytes (93.5 % ± 2.75 %) and 7 % to oligodendrocytes (6.50 % ± 2.75 %) based on their RNA expression profile, and no significant differences in cell population distribution were discovered between the transplanted mouse groups (suppl. figure 6b).

We used DESeq2^60^ to analyze differential expression of genes between ST and HT and Ctrl hiPSC-derived astrocytes, as well as mice transplanted with these cells, with gender as a covariate when applicable. For astrocytes, genes with Benjamini-Hochberg adjusted p-value <0.05, and absolute fold change >=1 were analyzed for Gene Ontology (GO) term over-representation using clusterProfiler^61^. For transplanted mouse cortices, the number of reads from human origin was too low for reliable comparison of the groups, and thus only mouse genome of the brains transplanted with hiPSC-astrocyte progenitors was further analyzed. We used genes with nominal p-value <0.05 to explore functional terms of the most significantly different genes. In addition, we analyzed these genes with knowledge-based Ingenuity Pathway Analysis (IPA; Qiagen). Heatmaps to compare sample disease and function pathways with the limited queries to neuronal tissues and CNS cell lines were drawn with R package gplots.

### Transplantation

hiPSC-glial progenitor cells from two female twin pairs (pair 1 and pair 5) and two sex-matched controls (control 2 and control 3) were selected for neonatal transplantation into Rag1 knockout mice forebrain. The protocol was adopted from Kim, et al. 2014^62^. Patient-derived hiPSC-glial progenitors from sphere culture were dissociated with Accutase (Stemcell technologies) and resuspended in the concentration of 100,000 cells/µl in PBS prior to transplantation. The neonatal pups were first cryo-anesthetized, and a total of 300,000 cells were injected into the brain at 5 locations: (0.8, 1.0, -1.5; 0.5μl); (−0.8, 1.0, -1.5; 0.5μl); (0.8, 2.0, -1.5; 0.5μl); (−0.8, 2.0, -1.5; 0.5μl); (0.0, -1.0, -1.5; 1μl). Altogether 155 pups were injected from 26 litters.

By the age of 10 months the transplanted cells had spread through the whole brain, including the olfactory bulb and cerebellum based on Hunu^+^ (human nucleus) cell spreading. Based on the spreading of positive cells for Hunu staining six mice were excluded from the results because of lack of transplantation success. There was mouse-to-mouse variation unrelated to particular mouse group in spreading and density of transplanted astrocyte progenitors based on human-specific GFAP, human-specific PDGFRα, and Transferrin (reactive for both human and mouse) (Figure 4b, suppl. figure 4 a-c) when scored blinded based on human GFAP and Hunu-immunoreactivities in standard regions of the brain. In ST mice there was a tendency towards lower density of human Hunu^+^ cells compared to Ctrl mice or HT mice (figure 4c-d).

### Immunohistochemistry

One half of the sacrificed mouse brain was fixed in 4% PFA, cryopreserved in sucrose, and frozen in OCT (Tissue-Tek OCT). Sagittal sections of 20 µm thickness were stained with the following primary antibodies: human nucleus, 1:250-1:400 (Mouse anti-HuNu, Millipore MAB4383), human-specific GFAP 1:500 (Mouse anti-GFAP STEM123 TakaraBio Y40420), human-specific PDGFRα, 1:300 (Rabbit anti-PDGF Receptor α, cat# 5241, Cell Signaling), and mouse/human reactive transferrin 1:500 (Rabbit anti-Transferrin Abcam, ab82411) Alexa Fluor goat anti-mouse (488 and 568, Invitrogen A11001 and A11004) and goat anti-rabbit (488 A11008) were used at 1:200 as secondary antibodies. The sections were embedded in Vector mounting medium for fluorescence with DAPI (H-1200).

### Animal handling

Animals were maintained in a controlled environment (temperature 22 ± 1°C and humidity 50–60%) on a 12h:12h light/dark cycle and provided with food and water *ad libitum*. Mice were studied in 2 cohorts: 1. cohort = 37 females, 2. cohort = 33 females (Total = 70 mice, Ctrl = 19, ST = 24, HT = 27). Truscan, plus maze, nesting, rotarod, NOR, social approach, odor ball, Y-maze and acoustic startle response were performed for 5- and 10-months old mice. Mice were weighed and handled before testing was started and moved to individual cages about one week before behavioral testing start. The experimenter was blind to groups when conducting the experiments. The experiments were conducted according to the Council of Europe (Directive 86/609) and Finnish guidelines and approved by the Animal Experiment Board of Finland.

### Activity (TruScan)

Motor activity of mice was automatically followed with infrared beam detection (TruScan, Coulbourn Instruments). Mice were tested for 10 min in dim light for different measured variables: move time, rest time, distance, stereotypy moves and time, vertical entries and time, jumps, ambulatory move time and distance.

### Elevated plus-maze

The mouse was placed onto the plus-formed elevated maze with two open and two closed arms (diameter 60 cm, center 5 cm x 5 cm, and arm width 5 cm). The mouse started from the center with the nose pointing to the closed arm, and was video-recorded for 5 min. Afterwards, the videos were analyzed with EthoVision XT 7 software to count the visits and the time spent in the open and closed arms.

### Nest building test

Nesting material and enrichement objects (plastic igloo) were removed from the home cages. One Nestlet cotton pad was placed in the cage overnight. After 19 h and 42 h the nests were photographed and scored on scale 0-4 based on the shape and coverage.

### Rotarod (Ugo Basile)

The mouse was placed on a rotating rod accelerating from 5 to 40 rpm in 5 min time. The latency to fall time and the spins on the rod were counted.

### NOR (Novel Object Recognition)

Two identical objects were placed symmetrically near the back corners of the cage, and the mouse was allowed to explore the objects for 20 min. Next day, the same objects were again put into the cage and the mouse behavior was video-recorded for 10 min. The time sniffing or otherwise in contact with the objects was measured by a stopwatch. If mouse did not explore the objects for at least 20 s, it was disqualified from the test. After a 1-h break a new session was run with one of the familiar objects replaced by a novel object. The exploration was video recorded for 5 min. The preference for the novel objects was calculated as follows: (exploration of novel object – exploration of familiar object) × 100% / total exploration time.

### Social Approach

The test was run in a three-compartment box. The empty middle compartment was connected with the lateral ones that both housed a cylindrical cage. A 4-month-old female mouse was placed inside one of the cages while the other one was left empty. The mouse was placed into the neutral central compartment and was video-recorded for 10 min. Social preference was calculated as follows: (exploration time in the mouse compartment – exploration time in the empty cage compartment) × 100% / total time in the lateral compartments.

### Odor ball –test

Every mouse was given one wooden ball (Ø 2 cm) to its home cage overnight. On test day 1, the mouse received new bedding and two balls, one impregnated with its own odor and another one from the cage of an unknown mouse. On test day 2, a ball impregnated with the mouse’s own odor was paired with a cardamom odorized ball. The position of the balls was randomized between mice. We measured the total sniffing time (the nose pointing to the ball at a distance < 2 cm) during a 120-s test session. Only if the total sniffing time exceeded 10 s, we calculated the odor preference as follows: (time sniffing unknown mouse/cardamom odor – time sniffing own odor) × 100% / total sniffing time.

### Spontaneous alternation in a Y-Maze

The mouse was put into the stem arm of a Y-shaped maze with black plastic walls and a transparent plastic ceiling to prevent peeking over the wall. If the mouse visited three different arms in a row, it got one alternation score. A total number of arm visits for the test was 20, and a maximum of 20 min was allowed for 10 last arm visits. The total number of alternations was counted through an overhead video monitor.

### Pre-pulse inhibition of acoustic startle

The acoustic startle response to the presentation of a sudden loud sound is a sensitive indicator of general reactivity of the animal. Pre-pulse inhibition, a reduction in the startle response to the sudden loud sound by a weak preceding sound pulse, is a sensitive measure of suppression of irrelevant stimulus from attention and is attenuated in patients with SCZ and several animal models of SCZ. The test was performed in a startle response system (TSE-Systems, Bad Homburg, Germany) that consisted of a sound-attenuated chamber (39 cm × 38 cm × 58 cm) and a cylindrical Plexiglas restrainer (8 cm × 8 cm × 25 cm) for the mouse located between two loudspeakers. The restrainer was located on a highly sensitive scale. The startle response was measured as apparent weight increase/decrease due to the muscle contractions. The mouse was first exposed for 5 min to a background white noise (65 dB) to adapt. The startle pulse (40 ms, 110 dB white noise) was presented alone preceded by a pre-pulse stimulus (20 ms tone) at three intensities (20, 35, 50 dB). The various pre-pulse tone + white noise combinations were presented in a pseudorandom order, 10 times each. The test duration was 35 minutes.

## Acknowledgements

We thank Marthe Van de Vliet, Eila Korhonen, Laila Kaskela, Sara Wojciechowski, Mirka Tikkanen and Velta Keksa-Goldsteine for technical help in generation and characterization of the stem cell lines.

## Author Contributions

J. Tiihonen and J. Koistinaho conceived the study. Š.L. planned and supervised differentiation of the astrocytes, cell line characterization, sample preparation for RNA sequencing, cell transplantation and histological stainings. M. Koskuvi differentiated the astrocytes, prepared RNA, FACS, experimental samples, run qRT-PCR, prepared co-cultures for calcium imaging, prepared RNA samples for sequencing and prepared the table for sex-specific DEGs. K.T., I. Hovatta performed RNA sequencing analyses and contributed to the interpretation of the results. T.C., J.L., S.T., J.S., and J. Kaprio gathered the data on twin pairs. I.O. and O.V. performed skin biopsies and rating of symptoms. M.L. contributed to the interpretation of the results. I. Hyötyläinen grew and differentiated astrocytes and helped in cell preparations. R. Giniatullina, R. Giniatullin, J. Tohka planned calcium-imaging studies together with M. Koskuvi, P.L.J.V., A.L. and C.R. who performed the calcium-imaging experiments and N.R., A.L. their analysis. M. Keuters transplanted the cells and together with H.D. perfused and collected tissue samples; L.P. prepared tissue samples for stainings and S.K. prepared for RNA and imaged brain sections. Y.C.W. imaged the whole-section scans and prepared the graphic figures based on them. H.T. planned and H.K. performed the behavior tests and statistical analyses; L.P. helped to perform the behavior tests. M. Koskuvi wrote the first draft of the manuscript and prepared the figures and tables with the help of Š.L., J. Tiihonen and J. Koistinaho.

