## Supplementary Information for "Patient iPSC-astrocytes show transcriptional and functional dysregulation in schizophrenia"

**Supplementary Table 1.** Summary of schizophrenia patients, their co-twins, healthy control participants, and hiPS lines used in this study.

**Supplementary Table 2.** Number of GO terms and IPA canonical pathways.

**Supplementary Table 3.** Synaptic transmission highlighting shared genes with Windrem et al. 2017

**Supplementary Table 4.** Statistics for calcium imaging results in glutamate application.

**Supplementary Table 5.** Statistics for calcium imaging results in GABA application.

**Supplementary Table 6.** Number of novel object recognition test excluded mice in each group.

**Supplementary Table 7.** Number of DEGs aligned with mouse genome.

**Supplementary Figure 1.** hiPSC-astrocyte characterization.

**Supplementary Figure 2.** IPA results for ST vs Ctrl comparison.

**Supplementary Figure 3.** Proportions of hiPSC-neurons in neuron-astrocyte cocultures.

**Supplementary Figure 4.** Transplanted Hunu+ cell characterization.

**Supplementary Figure 5.** Additional behavior test results.

**Supplementary Figure 6.** Transplanted mouse transcriptomic.

**Supplementary Table 1. Summary of schizophrenia patients, their co-twins, healthy control participants, and hiPS lines used in this study.** Pair 2 was excluded in the previous study (Tiihonen, et al. 2019).

| | Abbreviations of hiPSC lines | Group | Age at biopsy (years) | Sex | Medication |
|---|---|---|---|---|---|
| Pair 1 | SZ1 (HT 1) | Unaffected co-twin | 47 | Female | - |
| | SZ2 (ST 1) | Affected twin | 47 | Female | Clozapine |
| Pair 3 | SZ5 (ST 3) | Affected twin | 66 | Male | Zuclopenthixol |
| | SZ6 (HT 3) | Unaffected co-twin | 66 | Male | - |
| Pair 4 | SZ7 (ST 4) | Affected twin | 69 | Female | Previously clozapine, now sertindole and quetiapine |
| | SZ8 (HT 4) | Unaffected co-twin | 69 | Female | - |
| Pair 5 | SZ9 (HT 5) | Unaffected co-twin | 45 | Female | - |
| | SZ10 (ST 5) | Affected twin | 45 | Female | Clozapine |
| | SZ11 (Ctrl 1) | Control | 44 | Male | - |
| | SZ12 (Ctrl 2) | Control | 59 | Female | - |
| | SZ13 (Ctrl 3) | Control | 49 | Female | - |
| | SZ14 (Ctrl 4) | Control | 64 | Female | - |
| Pair 6 | SZ15 (ST 6) | Affected twin | 40 | Male | Olanzapine and quetiapine |
| | SZ16 (HT 6) | Unaffected co-twin | 40 | Male | - |
| | SZ17 (Ctrl 5) | Control | 63 | Male | - |
| | SZ18 (Ctrl 6) | Control | 50 | Female | - |

**Supplementary Table 2. Number of GO terms and IPA canonical pathways.** GO analysis done with genes DEG p.adj <0.05, log2FC >|1|, significant pathways listed (adj.p<0.05). IPA analysis done with genes DEG p.adj <0.05, log2FC >|1|, significant pathways listed (-log(p-value)>1.3).

| | Comparison | No of significant GO terms | No of significant pathways |
|---|---|---|---|
| ALL | ST vs. Controls | 67 | 19 |
| Female | ST vs. Controls | 39 | 11 |
| Male | ST vs. Controls | 450 | 65 |
| ALL | HT vs. Controls | 0 | 2 |
| Female | HT vs. Controls | 544 | 31 |
| Male | HT vs. Controls | 934 | 62 |
| ALL | ST vs. HT | 30 | 5 |
| Female | ST vs. HT | 0 | 5 |
| Male | ST vs. HT | 353 | 19 |

**Supplementary Table 3. Synaptic transmission with shared genes with Windrem et al. 2017.** The pathway is categorized as Cell-To-Cell Signaling and Interaction, Nervous System Development and Function; synaptic transmission; Synaptic transmission. The shared genes between our data and Windrem data are shown in bold. Related to figure 2e.

| | | p-value | z-score | Molecules in pathways (shared genes in Windrem, et al. 2017 in bold) |
|---|---|---|---|---|
| ST vs. HT | All | n.s. | - | |
| | Female | n.s. | - | |
| | Male | $1.23 \times 10^{-5}$ | 0.45 | LGI1, PRNP, GRIN2A, CNR1, RIMS1, APOE, LYPD1, NRXN1 |
| ST vs. Ctrl | All | n.s. | - | |
| | Female | $3.68 \times 10^{-4}$ | - | GRIK2, CNTNAP2, CDH8, KCNJ6 |
| | Male | $4.31 \times 10^{-7}$ | -0.71 | ADRB2, KCNMA1, SLC6A6, RAPGEF4, GRIA1, NPY2R, ALK, NTF3, GABRB2, KCNA1, GRM3, DLGAP1, **KCNJ9**, HLA-A, SCN1A, HAP1, PTK2B, PRKCB, GAD2, SLC1A3, PCDHB5 |
| HT vs. Ctrl | All | n.s. | - | |
| | Female | $1.38 \times 10^{-7}$ | -1.26 | SLC6A6, UNC13A, GRIK2, DPP6, EPHB2, **EPHB1**, LRP8, KCNJ6, RIMBP2, PTPRD, DGKB, SHANK2, CNTNAP2, KCNQ3 |
| | Male | $1.74 \times 10^{-9}$ | 1.79 | ADRB2, LGI1, GAD1, WNT5A, NR3C2, NQO1, KCNA1, GRM3, NLGN1, **KCND2**, CNTNAP2, NRG3, GRM5, GRM8, ADCY8, PCDHB5, **FGF12**, SCN2B, TNFRSF1B, RAPGEF4, GRIK2, NPY2R, ALK, RIMS1, NTF3, FBXO2, PLAT, NRXN1, CAMK2A, LYNX1, DLGAP1, HLA-A, CAV2, PMCH, SCN1A, GPR176, PCDHB16, APOE, LYPD1, CDH8 |
| Twins vs. Ctrl | All | n.s. | - | |
| | Female | $1.57 \times 10^{-6}$ | -2.41 | UNC13A, RAPGEF4, ERC2, GRIK2, CLSTN2, NBEA, KCNJ6, PTPRD, DGKB, SHANK2, CNTNAP2, LYPD1, CDH8, KCNQ3 |
| | Male | $5.08 \times 10^{-6}$ | 2.14 | ADRB2, SLC6A6, WNT5A, KCNA1, GRM3, **KCNJ9**, **KCND2**, CNTNAP2, GRM5, ADCY8, PCDHB5, SST, KCNMA1, TNFRSF1B, RAPGEF4, GRIK2, NPY2R, ALK, NTF3, CAMK2A, DLGAP1, HLA-A, PMCH, SCN1A, PCDHB16, CDH8 |

**Supplementary Table 4. Statistics for calcium imaging results in glutamate application.** Results from Tukey's multiple comparisons test. Related to figure 3d.

| **Glutamate** | **Mean Diff.** | **95.00% CI of diff.** | **Adjusted P Value** |
|---|---|---|---|
| **Male participants** | | | |
| Ctrl vs. HT | 0.02989 | -0.008615 to 0.06839 | 0.3166 |
| Ctrl vs. ST | -0.06296 | -0.09453 to -0.03140 | <0.0001 |
| HT vs. ST | -0.09285 | -0.1334 to -0.05233 | <0.0001 |
| Ctrl clozapine vs. HT clozapine | -0.02638 | -0.08053 to 0.02777 | 0.9123 |
| Ctrl clozapine vs. ST clozapine | 0.009666 | -0.03874 to 0.05807 | >0.9999 |
| HT clozapine vs. ST clozapine | 0.03605 | -0.007407 to 0.07950 | 0.2205 |
| **Female participants** | | | |
| Ctrl vs. HT | 0.02842 | -0.008840 to 0.06568 | 0.3442 |
| Ctrl vs. ST | -0.09674 | -0.1366 to -0.05688 | <0.0001 |
| HT vs. ST | -0.1252 | -0.1624 to -0.08794 | <0.0001 |
| Ctrl clozapine vs. HT clozapine | -0.06193 | -0.1045 to -0.01936 | 0.0001 |
| Ctrl clozapine vs. ST clozapine | 0.008534 | -0.02713 to 0.04419 | 0.9998 |
| HT clozapine vs. ST clozapine | 0.07047 | 0.02323 to 0.1177 | <0.0001 |
| **Sex differences** | | | |
| Ctrl male vs. Ctrl female | -0.06924 | -0.1041 to -0.03439 | <0.0001 |
| HT male vs. HT female | -0.0707 | -0.1114 to -0.03000 | <0.0001 |
| ST male vs. ST female | -0.103 | -0.1400 to -0.06599 | <0.0001 |
| Ctrl clozapine male vs. Ctrl clozapine female | -0.03224 | -0.07828 to 0.01380 | 0.4846 |
| HT clozapine male vs. HT clozapine female | -0.06779 | -0.1190 to -0.01656 | 0.0009 |
| ST clozapine male vs. ST clozapine female | -0.03337 | -0.07204 to 0.005291 | 0.171 |
| **Clozapine treatment** | | | |
| **Male participants** | | | |
| Ctrl vs. Ctrl clozapine | 0.04759 | 0.001630 to 0.09354 | 0.0347 |
| HT vs. HT clozapine | -0.00868 | -0.05667 to 0.03931 | >0.9999 |
| ST vs. ST clozapine | 0.1202 | 0.08518 to 0.1552 | <0.0001 |
| **Female participants** | | | |
| Ctrl vs. Ctrl clozapine | 0.08458 | 0.04963 to 0.1195 | <0.0001 |
| HT vs. HT clozapine | -0.00577 | -0.05026 to 0.03872 | >0.9999 |
| ST vs. ST clozapine | 0.1899 | 0.1494 to 0.2303 | <0.0001 |

**Supplementary Table 5. Statistics for calcium imaging results in glutamate application.** Results from Tukey's multiple comparisons test. Related to figure 3e.

| GABA | Mean Diff. | 95.00% CI of diff. | Adjusted P Value |
|---|---|---|---|
| **Male participants** | | | |
| Ctrl vs. HT | -0.002754 | -0.01117 to 0.005661 | 0.9959 |
| Ctrl vs. ST | -0.02191 | -0.02881 to -0.01502 | <0.0001 |
| HT vs. ST | -0.01916 | -0.02802 to -0.01030 | <0.0001 |
| Ctrl clozapine vs. HT clozapine | -0.01434 | -0.02618 to -0.002506 | 0.0043 |
| Ctrl clozapine vs. ST clozapine | -0.02033 | -0.03091 to -0.009748 | <0.0001 |
| HT clozapine vs. ST clozapine | -0.005986 | -0.01548 to 0.003512 | 0.6517 |
| **Female participants** | | | |
| Ctrl vs. HT | -0.01822 | -0.02636 to -0.01007 | <0.0001 |
| Ctrl vs. ST | -0.00587 | -0.01458 to 0.002842 | 0.5481 |
| HT vs. ST | 0.01235 | 0.004210 to 0.02048 | <0.0001 |
| Ctrl clozapine vs. HT clozapine | -0.004434 | -0.01374 to 0.004873 | 0.9242 |
| Ctrl clozapine vs. ST clozapine | 0.007298 | -0.0004964 to 0.01509 | 0.092 |
| HT clozapine vs. ST clozapine | 0.01173 | 0.001406 to 0.02206 | 0.0111 |
| **Sex differences** | | | |
| Ctrl male vs. Ctrl female | 0.01864 | 0.01102 to 0.02626 | <0.0001 |
| HT male vs. HT female | 0.003178 | -0.005718 to 0.01207 | 0.9913 |
| ST male vs. ST female | 0.03468 | 0.02659 to 0.04278 | <0.0001 |
| Ctrl clozapine male vs. Ctrl clozapine female | -0.004697 | -0.01476 to 0.005366 | 0.9338 |
| HT clozapine male vs. HT clozapine female | 0.005212 | -0.005988 to 0.01641 | 0.9352 |
| ST clozapine male vs. ST clozapine female | 0.02293 | 0.01448 to 0.03138 | <0.0001 |
| **Clozapine treatment** | | | |
| **Male participants** | | | |
| Ctrl vs. Ctrl clozapine | 0.02031 | 0.01026 to 0.03035 | <0.0001 |
| HT vs. HT clozapine | 0.008717 | -0.001772 to 0.01921 | 0.2181 |
| ST vs. ST clozapine | 0.02189 | 0.01423 to 0.02955 | <0.0001 |
| **Female participants** | | | |
| Ctrl vs. Ctrl clozapine | -0.004431 | -0.01194 to 0.003077 | 0.7412 |
| HT vs. HT clozapine | 0.01075 | 0.001026 to 0.02047 | 0.0159 |
| ST vs. ST clozapine | 0.01014 | 0.001290 to 0.01898 | 0.0099 |

**Supplementary Table 6. Number of NOR excluded mice in each group.** A high number of mice were excluded in NOR test because of their lack of attention to the objects. This was possibly caused by stress.

| | Ctrl group | | ST group | | HT group | |
|---|---|---|---|---|---|---|
| | 5 months | 10 months | 5 months | 10 months | 5 months | 10 months |
| All | 2/16 (12.5%) | 2/16 (12.5%) | 8/21 (38.1%) | 6/21 (28.6%) | 6/27 (22.2%) | 10/27 (37.0%) |
| RNAseq selected | 0/6 (0.0%) | 0/6 (0.0%) | 4/6 (66.7%) | 4/6 (66.7%) | 3/6 (50.0%) | 5/6 (83.3%) |

**Supplementary Table 7. Number of DEGs aligned with mouse genome.** The number of survived DEGs aligned with mouse genome at different cut-off values.

| | Mouse DEGs at different cutoffs: | | |
|---|---|---|---|
| **Comparison** | **Nominal p<0.05** | **P.adj<0.05** | **P.adj 0.05, absLFC>1** |
| **ST vs. HT** | **654** | **5** | **0** |
| **HT vs. Ctrl** | **362** | **0** | **0** |
| **ST vs. Ctrl** | **603** | **0** | **0** |
| **Ctrl vs. Non-transplant** | **2429** | **974** | **220** |

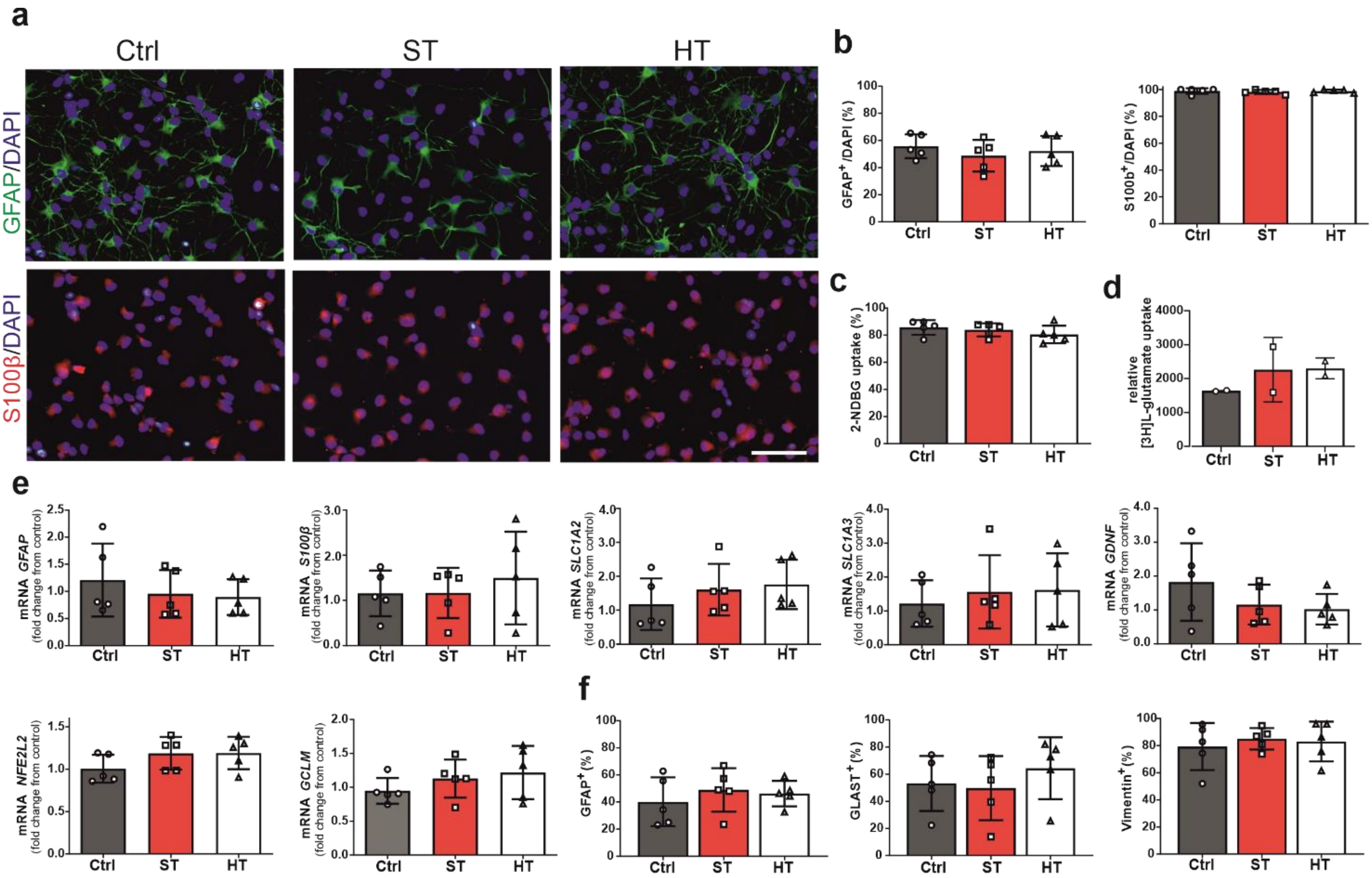

**Supplementary Figure 1. hiPSC-astrocyte characterization.** Related to Figure 1. All tested astrocytes exhibited typical stellate morphology and expressed *GFAP* (Ctrl: 55.7 % ± 8.8 %), *S100β* (Ctrl: 98.8 % ± 2.1 %), *GLAST* (Ctrl: 53.2 % ± 20.3 %), and *VIM* (Ctrl: 79.3 % ± 17.4 %). **a** The representative images of immunocytochemistry stainings for astrocytic markers GFAP (green) and S100β (red). Nuclei are counterstained with DAPI. Scale bar 50 µm. **b** Immunocytochemistry results showing the fraction of GFAP and S100β positive cells from each subject. **c** Glucose uptake analyzed by fluorescent glucose analog 2-NBDG and **d** glutamate uptake measured with radiolabeled glutamic acid. **e** The mRNA expression levels for astrocyte related genes GFAP, S100β, transporters SLC1A2 and SLC1A3, neurotrophic factor GDNF, and antioxidative genes NFE2L2 and GCLM measured by quantitative RT-PCR. **f** Flow cytometry detected proportions of GFAP, GLAST and Vimentin positive astrocytes. Ctrl=control subject, ST=affected twin, HT= unaffected co-twin. The data presented as mean ± SD, n=5 hiPSC lines, 1-2 replicates.

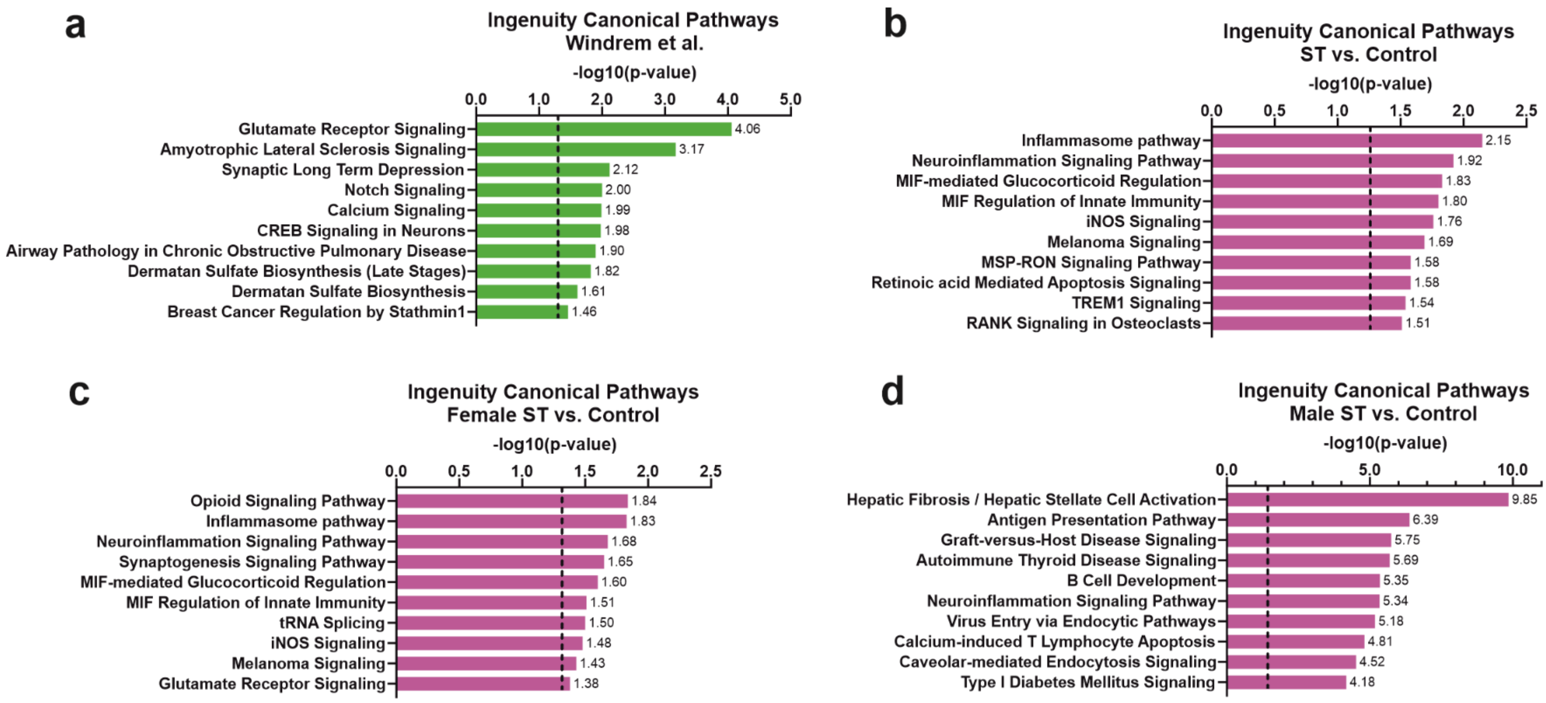

**Supplementary Figure 2. Ingenuity Pathway Analysis results for SCZ patients compared to control persons. a** The 10 most significant canonical pathways in the SCZ vs. Ctrl comparison from Windrem et al. (2017) data set. The pooled patient hiPSC-GPC gene expression (2 male and 2 female patients with childhood SCZ) was compared to the pooled gene expression pattern of hGPCs from 3 healthy control (2 males, 1 female). **b** The 10 most significant canonical pathways in ST vs. Ctrl comparison and separated **c** female and **d** male.

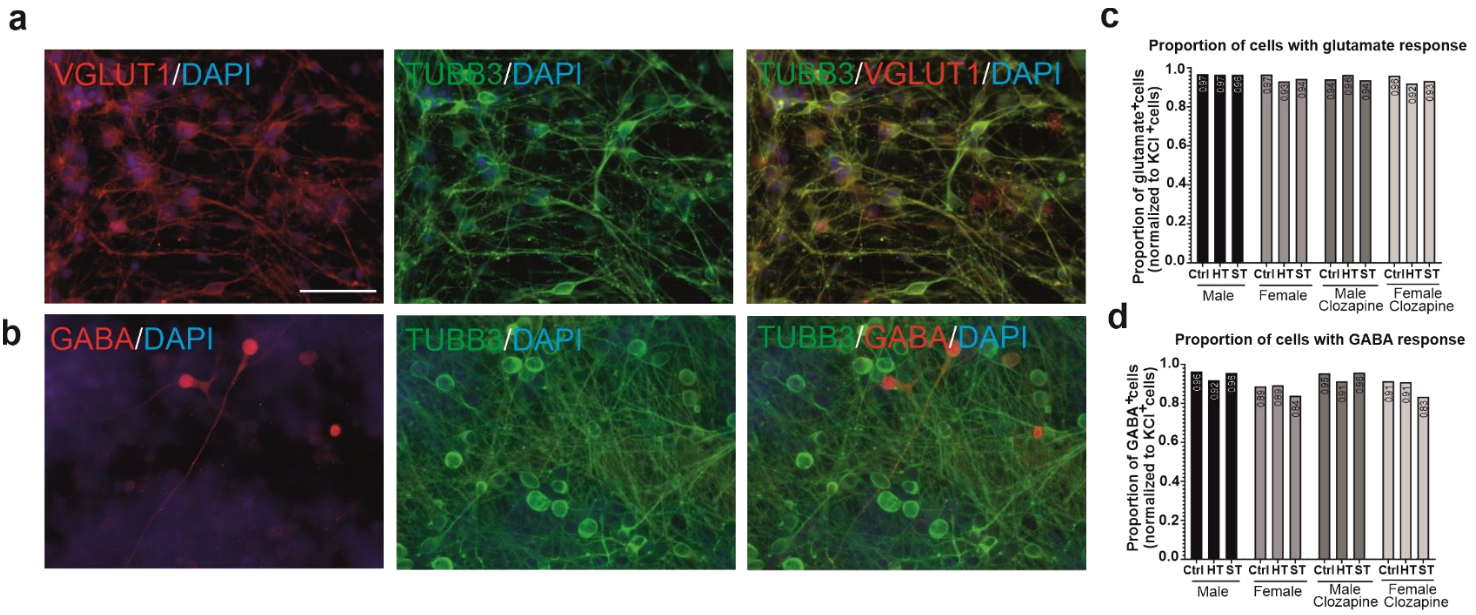

**Supplementary Figure 3. Proportions of hiPSC-neurons in neuron-astrocyte co-cultures.** Related to figure 3a. Representative pictures of **a** VGLUT1$^{+}$ and **b** GABA$^{+}$ neurons co-stained with TUBB3. The scale bar is 50 µm. Proportions of cells **c** responding to glutamate and **d** GABA according calcium recording. The bars present the proportion from pulled data. Regardless the group, from all the recorded cells (38,444) 46.9% responded to KCl (18,014 cells, threshold 5% of ΔF/F) and from the KCl responded cells 17,020 responding to glutamate (94.5%, threshold 0% of ΔF/F) and 15,893 cells to GABA (88.2%, threshold 0% of ΔF/F).

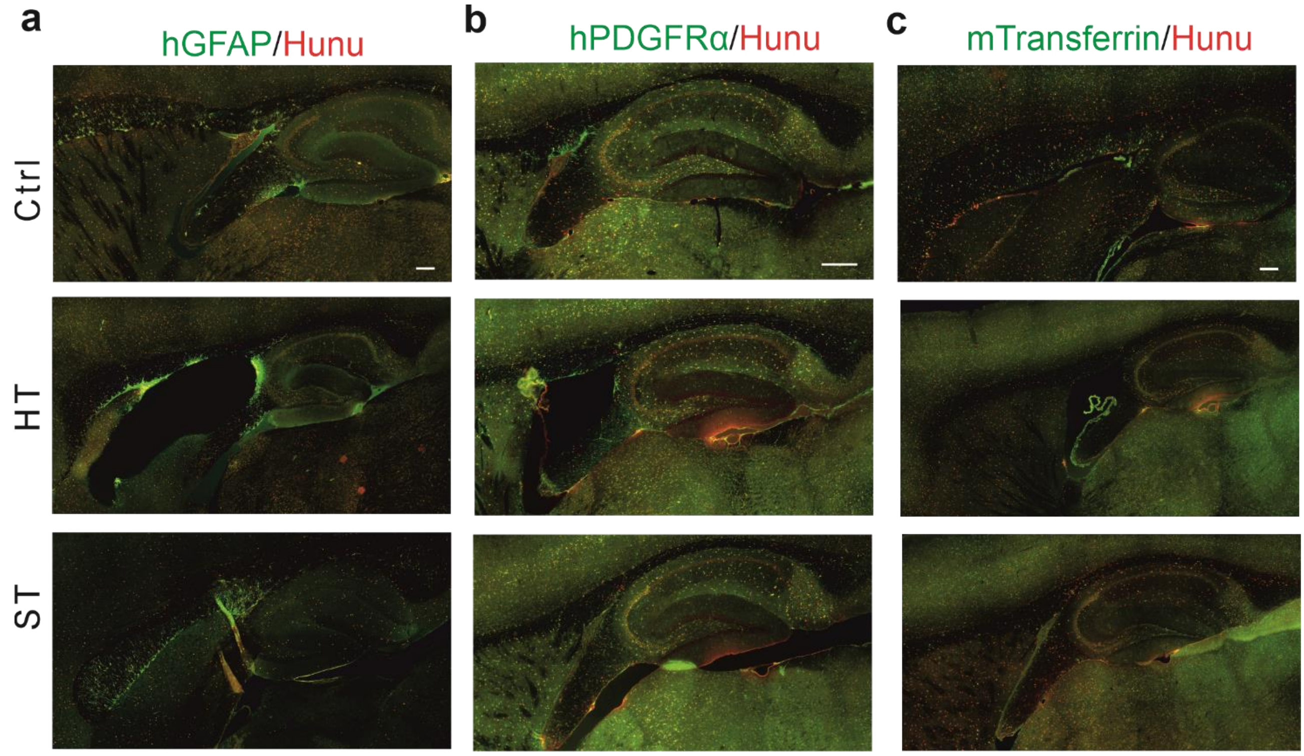

**Supplementary Figure 4. Transplanted Hunu⁺ cell characterization.** Related to figure 4b. Representative images of immuhistological stainings from **a** human-specific GFAP (green), **b** human-specific PDGFRα (green) and **c** mouse/human reactive Transferrin (green) stainings together with Hunu (human nuclei, red). Most of the Hunu+ cells showed positivity to glial progenitor marker PDGFRα and hardly any to oligodendrocyte marker Transferrin. Scale bar 200µm.

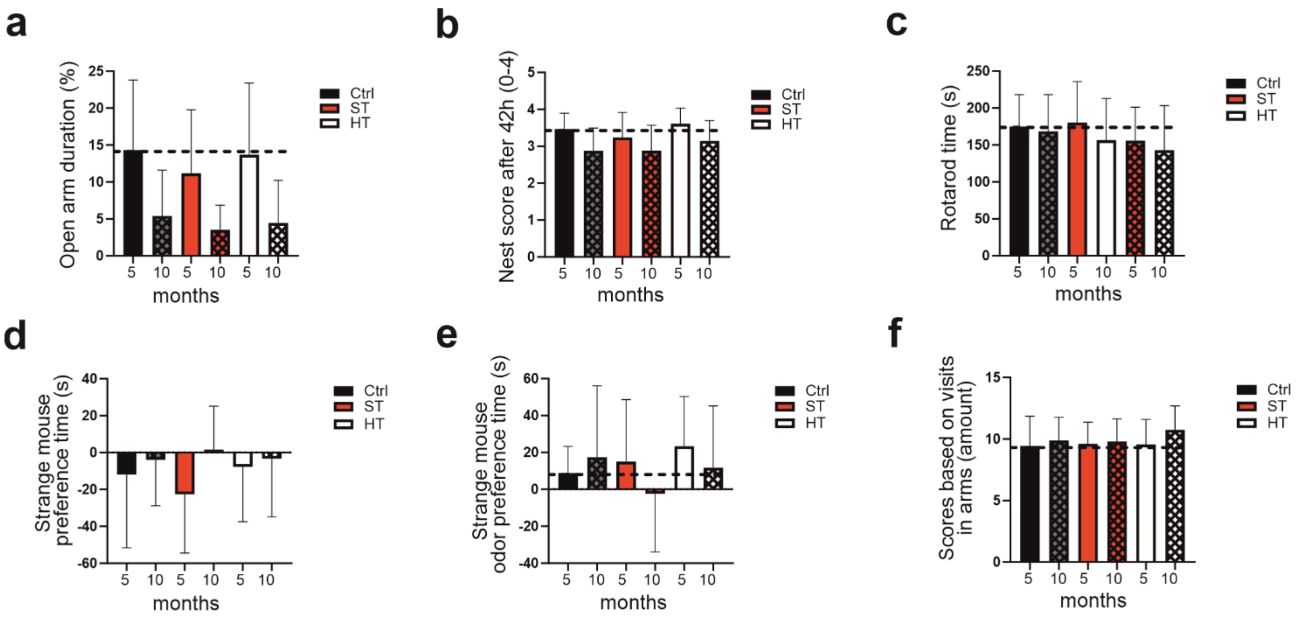

**Supplementary Figure 5. Additional behavior test results.** Related to Figure 4e. **a** Plusmaze showed no significant difference between treatment groups (p=0.14) but a highly significant age effect F(1.56) = 46.1, p < 0.001. **b** Nest building had no significant difference between treatment groups at 5 months (p = 0.39, N = 64, Kruskal-Wallis) or 10 months (p = 0.93). Significant age difference (p < 0.001, Wilcoxon Signed Ranks Test). Score range was 0-4 in nest building (4 best). **c** Mice were walking on accelerating rotating rod (max 360s). No significant difference between treatment groups (F(2.61) = 2.2, p = 0.12) in rotarod test and no age effect (F(1.61) = 2.5, p = 0.12). **d** Social approach had no difference between treatment groups (p = 0.83). Highly significant age effect (F(1.61) = 12.0, p < 0.001), but the age x treatment interaction did not reach significance (p = 0.15). **e** Sniffing preference time for unfamiliar mouse odor. No difference between the treatment groups (F(2.61) = 1.3, p = 0.28). **f** Alternations in Y-maze. No difference between the treatment groups (p = 0.37), age x treatment interaction p = 0.55. All data presented as Mean ± SD.

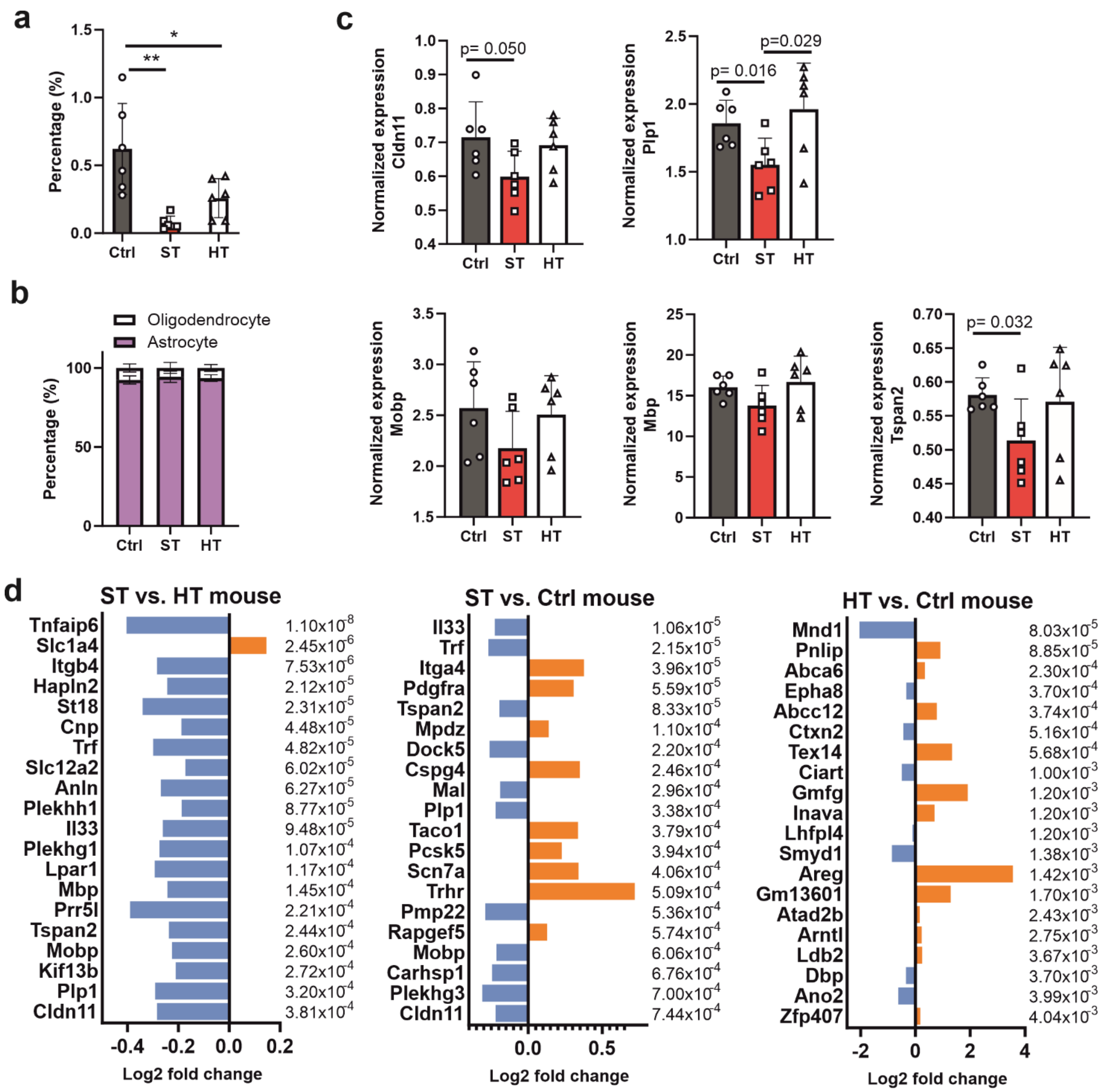

**Supplementary Figure 6. Transplanted mouse transcriptomic. a** Percentage of reads aligned in human genome. ** p < 0.01, * p < 0.05; 1-way ANOVA and Tukey's multiple comparison test. **b** Percentage of human cells in astrocyte and oligodendrocyte populations based on the human genome aligned transcriptomic profile. **c** Normalized expression of selected genes by RT-qPCR. Unpaired t-test. **d** The 20 most significant mouse DEGs in the ST vs. HT, the St vs. Ctrl and the HT vs. Ctrl comparisons in transplanted mice. All data presented as Mean ± SD.
